# Lancet2: Improved and accelerated somatic variant calling with joint multi-sample local assembly graphs

**DOI:** 10.1101/2025.02.18.638852

**Authors:** Rajeeva Lochan Musunuri, Bryan Zhu, Wayne E. Clarke, William Hooper, Timothy Chu, Jennifer Shelton, André Corvelo, Dickson Chung, Shreya Sundar, Adam M Novak, Benedict Paten, Michael C. Zody, Nicolas Robine, Giuseppe Narzisi

## Abstract

Here, we present Lancet2, an open-source somatic variant caller designed to improve detection of small variants in short-read sequencing data. Lancet2 introduces significant enhancements, including: 1) Improved variant discovery and genotyping through partial order multiple sequence alignment of assembled haplotype contigs and re-alignment of sample reads to the best supporting allele. 2) Optimized somatic variant scoring with Explainable Machine Learning models leading to better somatic filtering throughout the sensitivity scale. 3) Integration with Sequence Tube Map for enhanced visualization of variants with aligned sample reads in graph space. When benchmarked against enhanced two-tech truth sets generated using high-coverage short-read (Illumina) and long-read (Oxford Nanopore) data from four well characterized matched tumor/normal cell lines, Lancet2 outperformed other industry-leading tools in variant calling performance, especially for InDels. In addition, significant runtime performance improvements compared to Lancet1 (∼10x speedup and 50% less peak memory usage) and most other state of the art somatic variant callers (at least 2x speedup with 8 cores or more) make Lancet2 an ideal tool for accurate and efficient somatic variant calling.

## Introduction

Accurate and reliable detection of somatic Single Nucleotide Variants (SNVs) and insertions/deletions (InDels) is crucial for understanding the clonal structure and evolutionary dynamics of cancer. However, achieving comprehensive characterization of complex InDels remains a significant challenge for existing alignment-based somatic variant callers, largely due to reference bias ^1^. Lancet addressed this issue by performing local assembly and jointly analyzing tumor and normal data with colored de Bruijn graphs, thereby reducing reference bias and improving sensitivity for InDels in the 30–250 bp “twilight zone” ^2,3^.

Since its introduction, Lancet has become an integral part of the cancer genomics pipeline at the New York Genome Center ^4,5^ for liquid biopsy ^6^ and other efforts ^7–9^. Its effectiveness has also been demonstrated in a variety of independent studies across multiple cancer types including melanocytic naevi ^10^, polymorphous adenocarcinoma ^11,12^, pancreatic cancers ^11,13^, colorectal cancer ^14^, hyalinizing trabecular tumors of the thyroid ^15^, and triple-negative breast cancer ^16^.

Despite its success and increasing adoption, several limitations remain in the original Lancet framework. These included relatively long runtimes, high memory usage, and the absence of an interactive visualization tool to examine somatic variants with read support in graph space. To address these shortcomings, we developed Lancet2 — a redesigned and enhanced version of Lancet that offers improved performance, new genotyping strategy, and advanced somatic variant scoring with custom glassbox boosting models using InterpretML ^17–19^.

In this work, we compare Lancet2 against Lancet1 ^2^ and several industry-leading somatic variant callers, including Mutect2 ^20^, Strelka2 ^21^, DeepSomatic ^22^, and VarNet ^23^. To ensure thorough benchmarking, we developed enhanced “two-tech” truth sets for four widely used cancer cell lines — HCC1187, HCC1143, COLO829, and HCC1395 ^4,5^ — by integrating both high-coverage short-read Illumina and long-read Oxford Nanopore sequencing data. Our results demonstrate that Lancet2 not only improves variant calling accuracy, especially for InDels, but also significantly reduces runtime and memory usage while offering intuitive visualization tools to facilitate variant interpretation.

## Results

### Outline of the Lancet2 workflow

Lancet2 builds upon the joint multi-sample colored de Bruijn graph assembly strategy used in Lancet ^2^, introducing new modules for variant discovery, genotyping, (Figure 1) and filtering. Similar to the Lancet ^2^ tool, local haplotype contigs are generated by joint colored de Bruijn assembly of tumor and normal sample k-mers, through a process of graph clean-up, optimal k-mer size selection and Edmond-Karp style path traversal. To identify variants, Lancet2 performs Partial Order Alignment (POA)-based Multiple Sequence Alignment (MSA) of locally assembled contigs alongside the corresponding local reference sequence. Each sample read is subsequently re-aligned to the resulting MSA, ensuring that allele support is accurately determined by selecting the best-supported alleles for each read.

**Figure 1:**
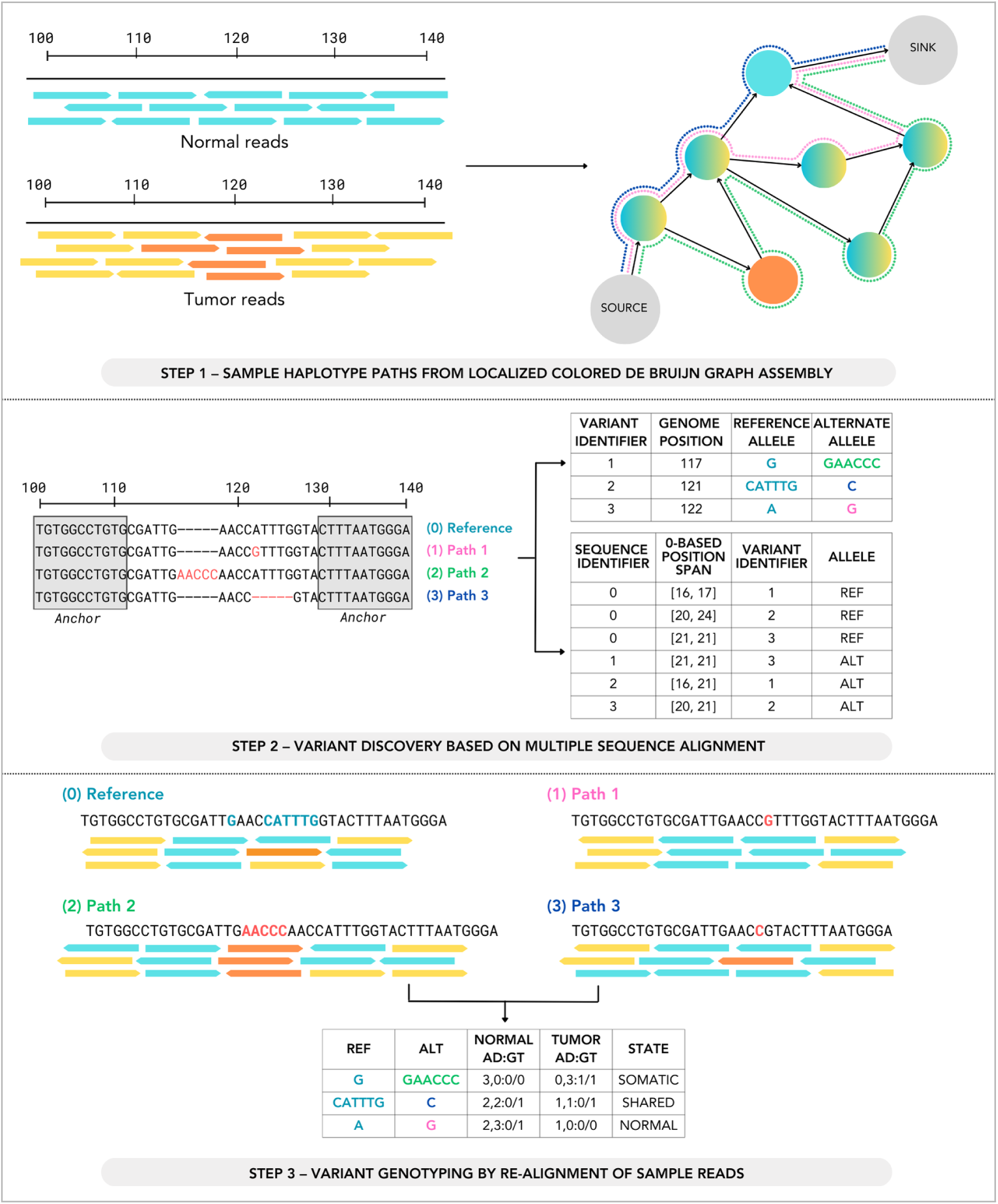
Illustration of various steps in the Lancet2 variant calling workflow. Step 1 – Representation of tumor and normal reads aligned to a localized colored de Bruijn graph, showing sample haplotype paths derived from the graph’s structure. Step 2 – Identification of candidate variants by performing partial order alignment–based multiple sequence alignments of assembled contigs and the reference. Variants are listed with their genomic positions, reference alleles, and alternate alleles. Step 3 – Genotyping of identified variants through the realignment of sample reads to the multiple sequence alignment, displaying normal and tumor allele depths and genotypes for each variant path.

Following variant discovery and genotyping, candidate variants are then scored and filtered using an interpretable, machine learning–driven glassbox boosting model. This model was trained on the candidate Lancet2 variant calls (including germline and mosaic artifacts) annotated with high confidence somatic truth set calls for the HCC1395 cell line ^4,5^ (Methods Section 2) using the InterpretML framework ^17–19^. Unlike opaque black-box models, this approach enables researchers to easily comprehend why specific variants are classified as somatic, providing both fine-grained explanations at the individual variant level and broader insights into the model’s overall behavior (Supplementary Figures 1 and 2).

### Enhanced Two-Tech Truth Sets for Cancer Cell Lines

We integrated high-coverage short-read (Illumina) and long-read (Oxford Nanopore) data (Table 1) to enhance previously published benchmarking truth sets for four well characterized cancer cell lines — HCC1187, HCC1143, COLO829, and HCC1395 ^4,5^. Beyond identifying variants supported by both platforms (COMMON), we systematically “rescued” variants initially observed by only one technology by examining raw alignments from the other platform (Supplementary Figure 3). This approach yielded three additional variant categories (Figure 2): variants called in long-read data and confirmed in short-read alignments (LR_ORIGIN), variants called in Illumina data and confirmed in long-read alignments (ILMN_ORIGIN), and variants without cross-platform evidence (DROPPED). Due to the lower sequencing coverage of the long read datasets, some of the low variant allele frequency calls in Illumina are not likely to be confirmed by long read data. However, variants called/missed in short/long read data due to technology specific mapping artifacts are greatly reduced as a result of this approach (Supplementary Figures 8 to 23). The resulting “two-tech” truth sets represent a robust resource for community-wide benchmarking of somatic SNV and InDel variant callers.

**Figure 2:**
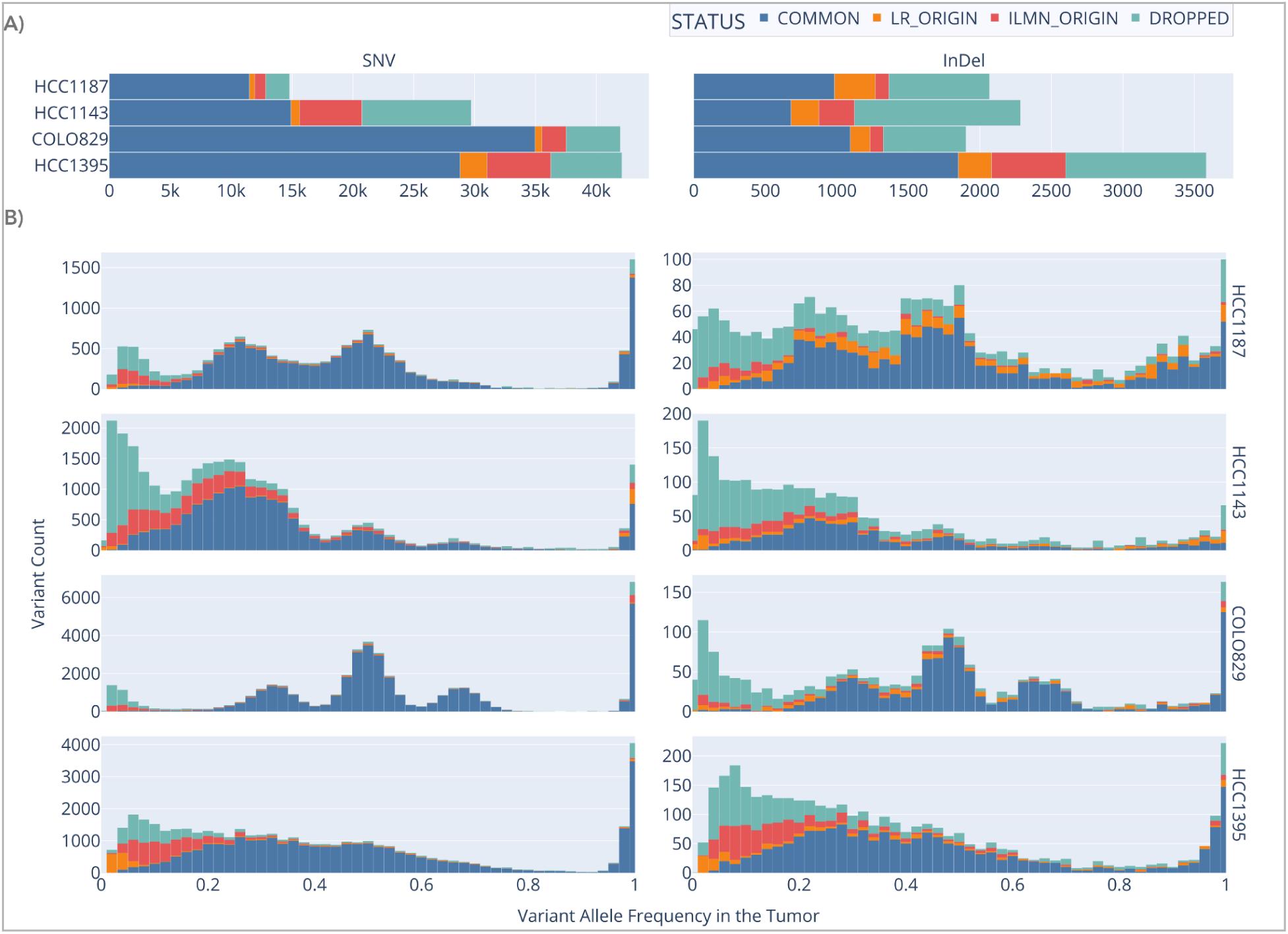
Visualization of the enhanced “two-tech” truth sets for four cancer cell lines (HCC1187, HCC1143, COLO829, HCC1395), showing variant classifications and variant allele frequency (VAF) distributions. (A) Stacked bar plots illustrating the counts of variants categorized as COMMON, LR_ORIGIN, ILMN_ORIGIN, and DROPPED for each cell line, grouped by variant type (Combined, SNV, InDel). (B) Histograms displaying the VAF distributions for each variant category and type, organized by cell line. Each row corresponds to a specific cell line, and each column presents the distribution of variants for SNV, or InDel categories. Short read data is used to extract the VAF for COMMON, ILMN_ORIGIN and DROPPED variants. Long read data is used to extract the VAF for LR_ORIGIN variants.

**Table 1:** Details of four cancer cell lines, including tumor and normal sample origins, sequencing coverage, and variant call sets derived from short-read Illumina NovaSeq 6000 data and long-read Oxford Nanopore R10.4.1 data. Each row provides information for one cell line, listing the tumor type, normal sample origin, and corresponding coverage values for both sequencing technologies.

| Cell Line | Tumor and Normal Origin | Short Read Data and Variant Calls |  |  | Long Read Data and Variant Calls |  |  |
| --- | --- | --- | --- | --- | --- | --- | --- |
| HCC1187 | Breast Ductal Carcinoma | Illumina NovaSeq 6000 | 103X | NYGC finalSet | Oxford Nanopore R10.4.1 | 101X | clairS |
|  | B-Lymphocyte-derived Normal |  | 70X |  |  | 41X |  |
| HCC1143 | Breast Ductal Carcinoma | Illumina NovaSeq 6000 | 315X | NYGC finalSet | Oxford Nanopore R10.4.1 | 118X | clairS |
|  | B-Lymphocyte-derived Normal |  | 178X |  |  | 34X |  |
| COLO829 | Metastatic Melanoma | Illumina NovaSeq 6000 | 255X | NYGC finalSet | Oxford Nanopore R10.4.1 | 120X | clairS |
|  | B-Lymphocyte-derived Normal |  | 203X |  |  | 38X |  |
| HCC1395 | Triple Negative Breast Cancer | Illumina NovaSeq 6000 | 85X | SEQC2 superSet in WGS NS_T_1 | Oxford Nanopore R10.4.1 | 106X | clairS |
|  | B-Lymphocyte-derived Normal |  | 71X |  |  | 38X |  |

### Variant Calling Performance Evaluation

Lancet2 achieves consistently higher InDel precision and F1 scores than competing variant callers across four distinct cancer cell lines (Figure 3). However for SNVs, DeepSomatic^22^ outperforms Lancet2 in precision and F1 score when evaluated using our enhanced “two-tech” truth sets (Figure 3). The default somatic model thresholds for Lancet2 are set at 95% probability for InDels and 90% for SNVs to ensure improved precision with a small impact on sensitivity. Users can fine-tune these cutoffs to prioritize higher sensitivity if their specific study demands it.

**Figure 3:**
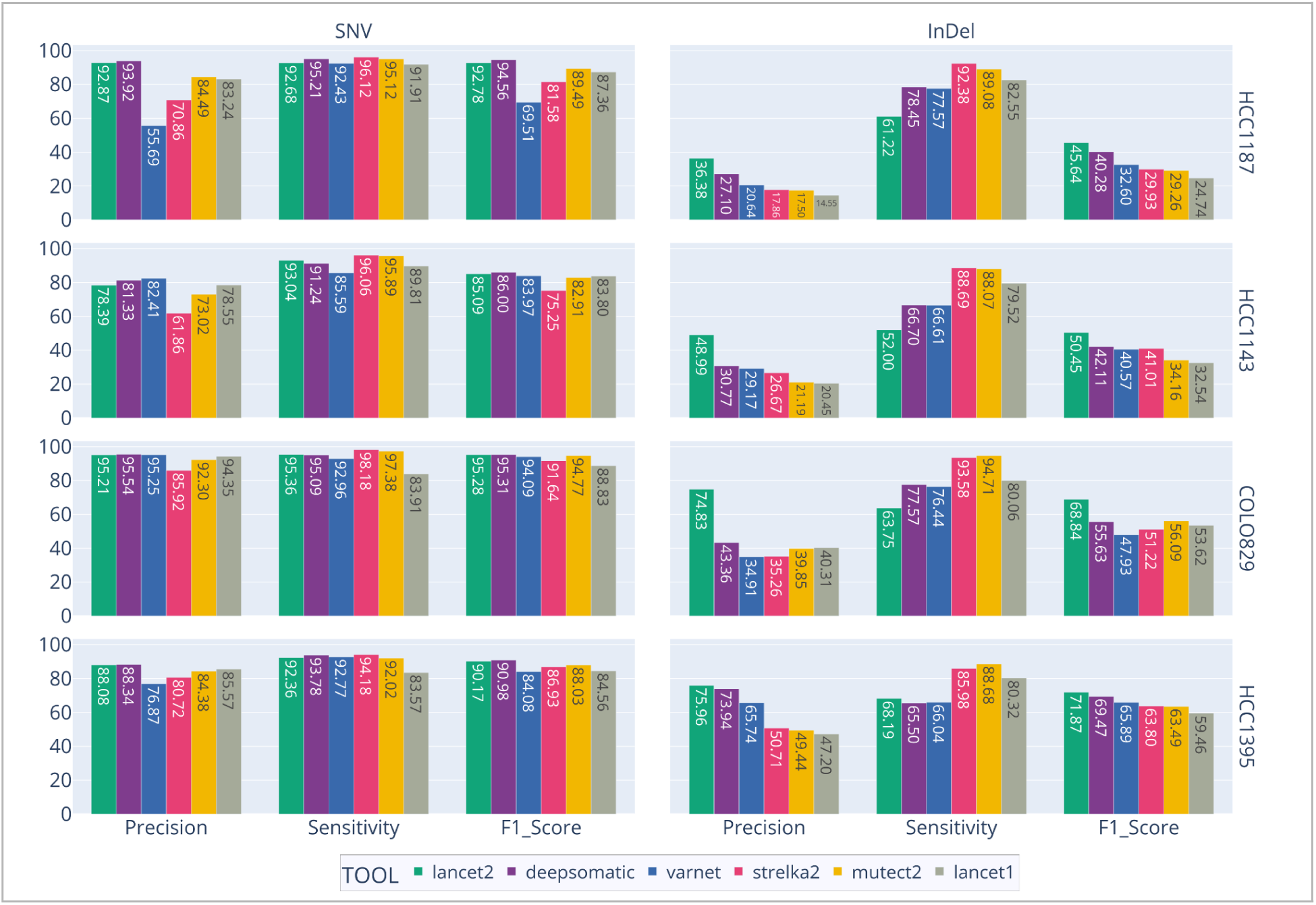
Comparison of variant calling performance metrics (precision, sensitivity, and F1 score) for multiple tools (Lancet2, DeepSomatic, VarNet, Strelka2, Mutect2, Lancet1) benchmarked against the enhanced “two-tech” truth sets. Each panel represents a distinct variant type (SNV, InDel) and each row corresponds to one of four cancer cell lines (HCC1187, HCC1143, COLO829, HCC1395). Bars indicate performance metric values for each tool, with displayed numerical labels.

Notably, Lancet2 maintains its performance advantage when benchmarked against previously published high-confidence truth sets ^4,5^, reinforcing the robustness of its improvements (Supplementary Figure 4). Moreover, this accuracy advantage extends across the full Variant Allele Frequency (VAF) spectrum, particularly for InDels. In COLO829, for example, other tools’ InDel calls suffer from elevated false-positive rates even at high VAFs, skewing their VAF profiles (Supplementary Figure 5). By contrast, Lancet2 consistently delivers accurate InDel calls, underscoring its robust and reliable performance in diverse benchmarking scenarios.

### Runtime Performance Evaluation

Lancet2’s local assembly approach inherently increases computational demands compared to hybrid or alignment-based methods. Nonetheless, Lancet2 achieves approximately a 10-fold speed improvement over the original Lancet, and in certain configurations, surpasses the performance of other tools due to its near-ideal scalability with additional CPU cores (Figure 4). Memory usage is similarly optimized, showing about a 40% reduction compared to Lancet1. This improvement becomes especially evident at higher coverage levels and when scaling to more cores (Supplementary Figure 6). These results highlight Lancet2’s ability to leverage CPU resources efficiently across a range of sequencing coverages and hardware configurations while maintaining low memory to CPU ratios.

**Figure 4:**
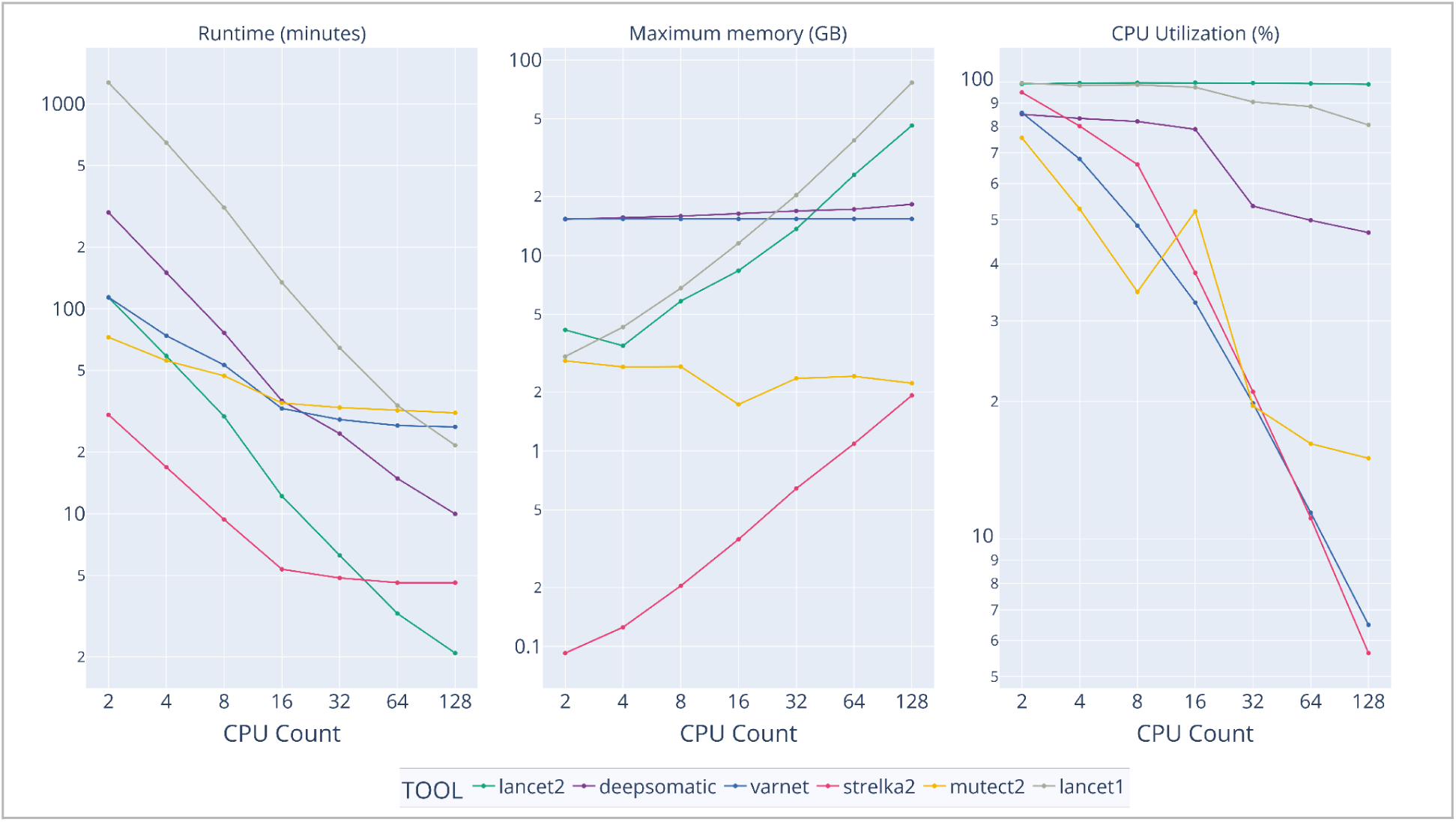
Runtime Performance by Number of CPUs: Benchmarked on Google Cloud with n2-highmem machines on chr20 HCC1395 vs HCC1395BL both at 64x coverage. Three line plots showing runtime in minutes, maximum memory usage in GB, and percent CPU utilization for multiple variant calling tools (Lancet2, Lancet1, DeepSomatic, Mutect2, Strelka2, VarNet) as the number of CPU cores increases. The x-axes represent increasing CPU counts, and the y-axes represent the respective metric values. Each line corresponds to a different tool.

### Graph Visualization of Variants using Sequence Tube Map

Lancet2 addresses the longstanding challenge of visualizing complex variants along with supporting reads from multiple samples in graph space, by its integration with Sequence Tube Map ^24^. Users can easily export the multiple sequence alignments used for variant discovery as GFA-formatted graphs which are then directly loaded into the Sequence Tube Map environment (Supplementary Information). Once there, somatic variants and their supporting read alignments from multiple samples can be intuitively explored, enabling clinicians and researchers to confirm allele support, understand local genomic complexity, and gain insights into variant structures (Figure 5 and Supplementary Figure 7). This interoperability provides a user-friendly, interactive web interface for examining detected variants at a granular level, facilitating improved interpretation, verification, and communication in both research and clinical contexts.

**Figure 5:**
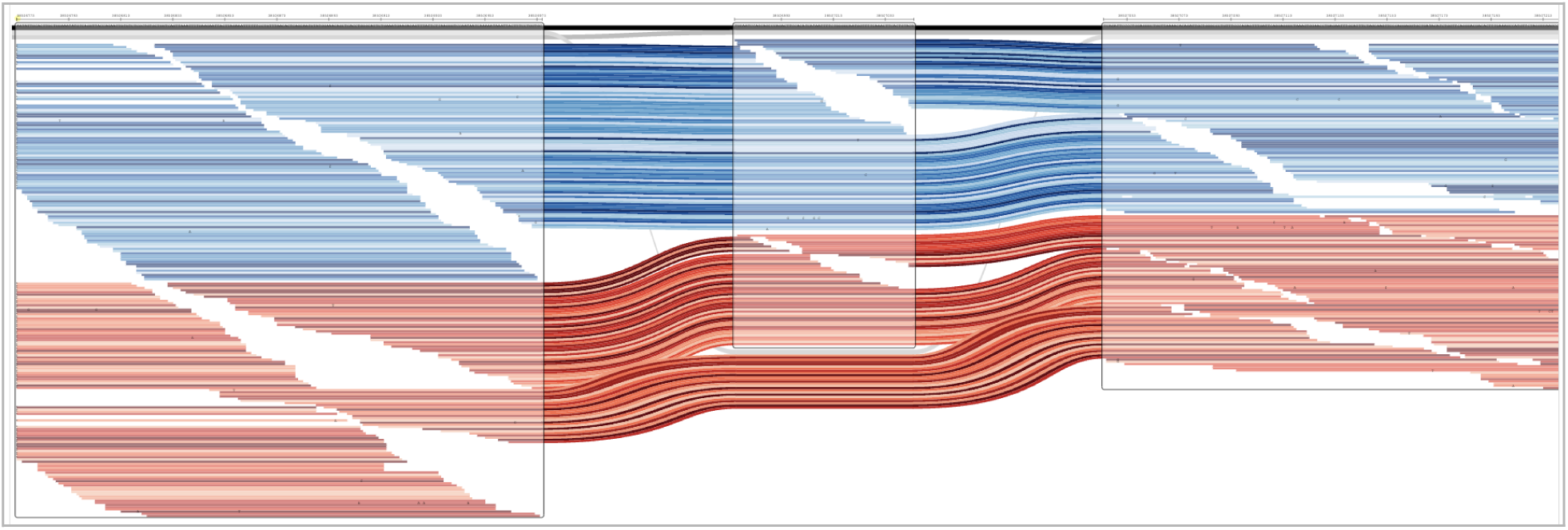
Sequence Tube Map visualization of a 70-bp somatic heterozygous deletion in HCC1395 tumor and normal cell lines with sample reads aligned to the local Lancet graph. The blue tracks correspond to normal sample reads, the red tracks below them correspond to tumor sample reads, and the grey boxes correspond to the nodes of the Lancet2’s exported sequence graph. The red tracks at the bottom that skip the middle box are the deletion reads in the tumor sample.

## Discussion

We present Lancet2, a next-generation somatic variant caller that significantly improves upon Lancet1 in accuracy, efficiency, interpretability, and visualization. Our benchmarking results show that Lancet2 outperforms other industry-leading tools in InDel variant calling performance, including Mutect2, Strelka2, VarNet, DeepSomatic, and the original Lancet, while maintaining competitive performance for SNVs. This aligns with previous studies highlighting the challenges of accurately identifying complex InDels using alignment-based approaches ^2,3^, emphasizing the value of local assembly methods in capturing subtle, structure-rich variants that may profoundly influence our understanding of tumor heterogeneity and clonal evolution.

A key advance in this work was the generation of enhanced benchmarking truth sets derived from integrating Illumina short-read and Oxford Nanopore long-read data for four commonly used cancer cell lines. By leveraging both sequencing platforms and “rescuing” variants identified by only one technology through cross-examination of raw alignments from the other, we produced a more robust “two-tech” truth set. One limitation of the two-tech approach and the resulting truth set is the inability to confirm low variant allele frequency variants due to the cost prohibitiveness of high depth long read sequencing. Nevertheless, this expanded resource represents a meaningful step forward, providing a more robust benchmark than purely short-read–based sets, and enabling a more realistic evaluation of variant calling performance. As long-read sequencing becomes more pervasive, such integrative approaches to truth set construction are increasingly recognized as critical for pushing the boundaries of somatic variant calling ^4,5^.

Beyond variant detection accuracy, Lancet2 demonstrates marked improvements in runtime and memory efficiency. Through careful algorithmic optimization and parallelization, Lancet2 is about 10 times faster and 40% more memory efficient than the original Lancet, making it more practical for large-scale and cloud-based analyses. Notably, Lancet2’s nearly linear scalability with increasing CPU cores and its relatively stable resource consumption at higher coverage levels set it apart from many existing tools. While previous studies have documented challenges in scaling variant callers to larger datasets and higher coverage ^20,21^, our findings indicate that resource-efficient local assembly is achievable and can be integrated into routine pipelines without compromising accuracy. In fact, Lancet2’s modular nature and its speed improvements allow users to run pangenome style – multiple tumor, multiple normal joint calling to generate raw candidate variants in joint VCF format. Future work is planned to simplify somatic variant analysis for longitudinal projects by integrating out of the box custom filtering and annotation when using multi-sample (more than 2 samples) mode.

The improved interpretability of Lancet2’s somatic filtering, achieved by incorporating a transparent, intelligible machine learning model ^19^, is another important milestone. The ability to explain classification decisions both locally (for individual variants) and globally (across the full model) stands in contrast to more opaque “black-box” filtering algorithms. This level of model transparency is increasingly valued in both research and clinical settings, where decision-making often requires careful scrutiny and validation of variant calls.

In addition, Lancet2 introduces graph-based visualization of cancer variants by its integration with Sequence Tube Map. This capability meets a pressing need to visualize both simple and complex variants and their supporting reads directly in graph space. Similar graph-centric approaches have been proposed in other genomic studies to better represent structural complexity and reduce reference bias ^25,26^, but their adoption in routine somatic variant interpretation has lagged behind. This integration allows researchers and clinicians to intuitively navigate complex genomic regions, enhancing transparency and confidence in variant calls — particularly those that challenge conventional linear-reference paradigms.

Despite these advances, Lancet2 currently focuses on small variants, and broader applicability to larger structural events will require future work. Additionally, generating comprehensive two-tech truth sets demanded extensive data, underscoring the need for ongoing community efforts to produce standardized, robust benchmarks. Furthermore, the adaptability of Lancet2’s filtering model to entirely new sequencing technologies or cancer types has yet to be comprehensively assessed and may require retraining or calibration as new data types emerge.

In summary, Lancet2 represents a substantial step forward in the accuracy, efficiency, and interpretability of somatic variant calling. Its performance improvements over industry-leading methods and its validated robustness across multiple well-characterized cancer cell lines suggest that it can serve as a valuable tool in both research and clinical genomics workflows.

## Methods

### 1. Variant discovery and genotyping workflow

Lancet2 builds on the previously described joint multi-sample Colored De-Bruijn Graph (cdbg) based assembly framework ^2^. As in the original Lancet approach, it processes the genome in consecutive, overlapping windows to perform read selection, graph construction, graph cleaning, active region detection, and path enumeration using an Edmond-Karp–style algorithm ^27^. Building on this foundation, Lancet2 introduces additional modules for variant discovery, genotyping, and scoring.

After assembling the local graph, Lancet2 aligns the assembled contigs and the corresponding reference sequence using a SIMD-accelerated partial order multiple sequence alignment (MSA) algorithm ^28–30^. Variants detected in these MSAs are recorded with their genomic coordinates, MSA positions, and reference/alternate alleles (Figure 1, Step 2).

Subsequently, all constituent reads used in assembly are re-aligned to both the local reference and assembled contigs with the minimap2 library ^31,32^. Read alignments are then sorted by descending gap-compressed identity and alignment score before they are assigned as support using exact sequence match to the discovered variant alleles, ensuring each read contributes only once per variant. These support counts form the basis for genotype determination and allele frequency estimation in the sample.

### 2. Building somatic glassbox boosting model

We trained the somatic glassbox boosting model using data from HCC1395 WGS_NS_T_1 (tumor) and WGS_NS_N_1 (normal) datasets. FASTQ files for both libraries were obtained from the Sequence Read Archive (SRA; ID:SRR7890893 for tumor and ID:SRR7890943 for normal) and processed through the NYGC cancer pipeline ^4^ to produce final alignments. Lancet2 was run on these tumor/normal pairs to generate a full set of raw candidate variants which include germline, mosaic and false positive artifacts (non-somatic) in addition to somatic variants. These calls were annotated against the previously published high-confidence truth set for HCC1395 ^5^, labeling them as true somatic or non-somatic. Chromosome 1 variants were excluded for later testing.

Each candidate variant was characterized by a number of metrics/features, including window-level (percent high quality reads per sample), variant-level (type, length, depth, absolute variant allele frequency difference between tumor and normal), and allele-level (allele counts, strand bias, base quality, mapping quality) features which are then utilized in the model training process. Due to an extreme class imbalance (∼1 true somatic per 200 non-somatic. See Supplementary Section 1 for more details), we applied random undersampling to reduce the ratio to about 1:30. After undersampling, we trained an Explainable Boosting Machine (EBM) using the InterpretML framework ^19^. The resulting trained model was saved to disk in Python’s native pickle format, enabling its direct integration into Lancet2’s filtering pipeline for somatic variant scoring. This integration greatly simplifies and improves the filtering process compared to multiple empirical hard cutoffs by relying on a single probabilistic score that can be used by the end user to balance precision and sensitivity.

### 3. Enhanced “two-tech” truth set generation

The “two-tech” truth sets integrate short-read (Illumina) and long-read (Oxford Nanopore) variant calls, leveraging cross-technology validation to improve confidence and accuracy. For each of the four selected cancer cell lines, we began with the full output of the NYGC cancer pipeline or the superSet calls ^4,5^ (short-read) and the PASS-filtered clairS v0.3.0 ^33^ variants (long-read), intentionally including low-confidence calls to allow subsequent cross-validation. These call sets are compared using RTG vcfeval ^34^ and the resulting true positive variants from the short-read call set are included in the “two-tech” truth set as the “common” variants.

Beyond identifying variants present in both callsets, we attempted to “rescue” uniquely detected variants by performing a pileup-based genotyping step using Freebayes ^35^ in pooled continuous mode. Variants exclusive to one platform’s calls were included in the final “two-tech” truth set only if the alternate allele was supported by at least two reads at ≥ 20× coverage in the other platform’s data. For the HCC1395 cell line, we excluded all variants from chromosome arms 6p, 16q and chrX because somatic and germline variants cannot be distinguished due to copy number loss in the normal sample ^5^. Final variant intersections between short/long calls and the Freebayes calls were conducted using RTG vcfeval ^34^, ensuring that only cross-validated variants contributed to the enhanced truth sets.

### 4. Architectural improvements

Lancet2 introduces several enhancements that streamline its architecture, improve interoperability, and increase performance. Following multiple sequence alignment (MSA), local assembly graphs can now be exported in GFA format, facilitating integration with a variety of external visualization and analysis tools (e.g., Bandage ^36^, VG ^37^, Sequence Tube Map ^24^). Direct integration with htslib ^38^ allows Lancet2 to accept both BAM and CRAM alignment files, offering users flexibility in balancing storage and analysis constraints. Adopting modern C++20 language features ensures improved maintainability and testability over time. In addition, Lancet2 employs a pull-based, reactive multithreading model with a fast, lock-free concurrent queue ^39^. This model dynamically distributes “work units” (genomic regions to process) across multiple worker threads, maximizing CPU utilization and optimizing performance throughout the entire runtime.

### 5. Features and Improvements to Sequence Tube Map

Several enhancements were introduced to Sequence Tube Map to support visualization of Lancet2 variants using the exported GFA graphs along with aligned supporting reads. First, the input data selection system was generalized to handle multiple read “tracks” from different sources, requiring internal refactoring, new user interface components, and in-app documentation. Sequence Tube Map can now load any number of distinct read tracks, along with graph or haplotype tracks, each displayed separately. This enables clear distinction between tumor and normal read sets from Lancet2 output. A new dialog provides per-read details, including graph traversal paths.

To streamline loading Lancet2 outputs, support for pre-defined visualizations (“chunks”) was extended to let chunks carry their own track configuration, similar to UCSC Genome Browser sessions ^40,41^. These chunks can be fetched from remote servers using documented, testable URL formats, and users can navigate among them easily. This workflow integrates end-to-end with Lancet2’s prep_stm_viz.sh script, producing interactive visualizations directly from BAM/VCF inputs (See “Lancet Example” in the Sequence Tube Map demo server – https://vgteam.github.io/sequenceTubeMap). The full end to end workflow showing this integration between Lancet2 and Sequence Tube Map is available in Supplementary Section 5.

Finally, visualization drawing algorithms were refined to accommodate higher coverage and multiple BAM files. Node spacing was adjusted to ensure adequate horizontal space for each read path, and the coordinate bar now supports breaks due to these expansions. Read ordering logic was also improved to handle multiple tracks under complex topologies, including inversions and cycles.

## Supporting information

somatic_EBM_model_features.txt

## Data and Code Availability

Lancet2 is an open source software distributed under a BSD 3-Clause License at https://github.com/nygenome/Lancet2.

Short read Illumina NovaSeq and Long read Oxford Nanopore PromethION alignment files for the four cancer cell lines and their matched normals used in the “two-tech” truth set generation process are available at the following SRA accession IDs.

Short read Illumina data –

COLO829 (SRX6743552), COLO829BL (SRX6743554),
HCC1187 (SRX6743560), HCC1187BL (SRX6743562),
HCC1143 (SRX6743556), HCC1143BL (SRX6743558),
HCC1395 (SRX4728475), HCC1395BL (SRX4728425)

Long read Oxford Nanopore data –

COLO829 (SRX27516919), COLO829BL (SRX27516920),
HCC1187 (SRX27516921), HCC1187BL (SRX27516922),
HCC1143 (SRX27516923), HCC1143BL (SRX27516924),
HCC1395 (SRX27516925), HCC1395BL (SRX27516926)

The resulting “two-tech” truth set variant call files (VCF) are available at the following publicly accessible google cloud storage paths –

gs://lancet2-paper/truths/COLO829/two-tech_truth/COLO829.TwoTechTruth.final.vcf.gz
gs://lancet2-paper/truths/HCC1187/two-tech_truth/HCC1187.TwoTechTruth.final.vcf.gz
gs://lancet2-paper/truths/HCC1143/two-tech_truth/HCC1143.TwoTechTruth.final.vcf.gz
gs://lancet2-paper/truths/HCC1395/two-tech_truth/HCC1395.TwoTechTruth.final.normal_loh_r egions_excluded.vcf.gz

## Supplementary Information

### 1. Somatic glassbox boosting model

Lancet2 v2.8.4-main-6ef7ba445a was used with default parameters to call all raw variants with alternate allele support greater than at least 2 reads from the HCC1395 vs HCC1395BL NovaSeq libraries obtained from SRA (SRR7890893 for tumor and SRR7890943 for normal) and processed through the NYGC cancer pipeline^4^. RTG vcfeval^34^ was then used to compare the raw Lancet2 variants with the previously published high-confidence truth set for HCC1395^5^, resulting in the classification of the raw variants into 7,798,060 false positive variants missing in the truth set and 39,249 true positive variants present in the truth set.

~~~
Lancet2 pipeline --num-threads 224 \
--reference GRCh38_full_analysis_set_plus_decoy_hla.fa \
--out-vcfgz HCC1395.Lancet_v2.8.4-main-6ef7ba445a.vcf.gz \
--normal HCC1395BL_SAMN10102574_SRR7890943.bam \
--tumor HCC1395_SAMN10102573_SRR7890893.bam
~~~

~~~
rtg vcfeval --output-mode=annotate --template \
GRCh38_full_analysis_set_plus_decoy_hla.sdf \
--all-records --vcf-score-field QUAL --sample=“ALT,ALT” \
--evaluation-regions SEQC2_High-Confidence_Regions_v1.2.bed \
--baseline “SEQC2.high-Confidence_Combined.v1.2.1.vcf.gz” \
--calls HCC1395.Lancet_v2.8.4-main-6ef7ba445a.vcf.gz \
--output rtg_vcfeval_output_HCC1395
~~~

All the variants from chromosome 1 were left out in the training process in order to be used in the later testing/benchmarking process. Due to the extreme class imbalance between the false vs true positive sets, the false positive variants were randomly undersampled to pick only one million variants. The full list of all the features that were extracted into a dataframe for 34,176 true positive somatic variants and 1,000,000 false positive non-somatic variants are available in Supplementary file 2.

Explainable Boosting Classifier from InterpretML^19^ v0.5.1 was then used to train the model with parameters max_bins=32, smoothing_rounds=2000, max_rounds=25000. The resulting somatic glassbox boosting model that was built is accessible publicly at https://storage.googleapis.com/lancet-ml-models/somatic_ebm.lancet_6ef7ba445a.v1.pkl.

Global term/feature importances of the somatic glassbox model can be explored (Supplementary Figure 1) using the following snippet of python code.

~~~
import pickle
import interpret
with open(“somatic_ebm.lancet_6ef7ba445a.v1.pkl”, “rb”) as rf:
     model = pickle.load(rf)
     interpret.show(model.explain_global())
~~~

Local explanation for why a single variant is classified as somatic/non-somatic can also be explored (Supplementary Figure 2) using the following python code snippet.

~~~
interpret.show(model.explain_local(X, Y))
~~~

This enables end users and researchers to provide a definitive answer to why a particular variant was marked as somatic or non-somatic by the machine learning model.

### 2. Enhanced Two-Tech Truth set generation

For the HCC1187, HCC1143 and COLO829 cell lines, previously published NYGC v6 cancer pipeline final calls ^4,5^ (includes lower confidence calls with support from a single variant caller as well) was used as the short read call set. For the HCC1395 cell line, v1.2 superSet calls from the Sequencing Quality Control Phase 2 Consortium ^4,5^ (SEQC2) were used as the short read call set.

Long read data for the 4 cancer cell lines were generated on the Oxford Nanopore R10 flow cell. Prior to variant calling, each sample dataset was base called using dorado v0.7.1_80da5f5 with the v4.1.0 super high accuracy (sup) model followed by read alignment to GRCh38 reference genome using minimap2 v2.28-r1209. PASS calls from clairS v0.3.0 were then used as the long read call set.

The clairS command line used to generate per chromosome variant calls for the long read call set is as follows –

~~~
/usr/bin/time --verbose /opt/bin/run_clairs \
    --tumor_bam_fn ∼{tumorCram} \
    --normal_bam_fn ∼{normalCram} \
    --ref_fn ∼{referenceFasta} \
    --platform ont_r10_dorado_sup_4khz \
    --enable_indel_calling --ctg_name ∼{contigName} \
    --sample_name ∼{tumorSampleName} --threads 64 \
    --output_dir “$(pwd)” \
    --output_prefix “∼{outFilePrefix}.snv” \
    --indel_output_prefix “∼{outFilePrefix}.indel”
~~~

In order to generate the “two-tech” truth set, RTG vcfeval v3.12.1 was used to intersect the short read and long read call sets. The RTG command line used to generate the intersection is shown below.

~~~
rtg vcfeval \
--template GRCh38_full_analysis_set_plus_decoy_hla.sdf \
--baseline ${SHORT_READ_TRUTH_SET_CALLS} \
--calls ${LONG_READ_CLAIRS_PASS_CALLS} \
--all-records --sample=ALT,ALT --vcf-score-field=QUAL \
--output-mode=split --output LR_vs_SR_output
~~~

The variants that are common between both the short and long read call sets form the initial set of “two-tech” callset. We then attempted to rescue the variants that are uniquely seen either in the short or long read call sets into the “two-tech” truth set by inspecting for evidence of variants in the raw alignments, using Freebayes v1.3.8 in pooled continuous mode. The command line that was used to inspect for evidence of variants in long or short read alignments is as follows –

~~~
# Attempt to rescue variants uniquely present
# in the short read callset into “two-tech” calls
# by looking for them in long read alignments
/usr/bin/time --verbose freebayes \
--fasta-reference GRCh38_full_analysis_set_plus_decoy_hla.fa \
--variant-input ${SHORT_READ_CALLSET_UNIQUE_VCF} \
--targets ${SHORT_READ_CALLSET_UNIQUE_REGIONS_BED} \
--only-use-input-alleles --hwe-priors-off \
--binomial-obs-priors-off --allele-balance-priors-off \
--no-population-priors --legacy-gls \
--pooled-continuous --pooled-discrete \
--min-alternate-fraction 0 --min-alternate-count 1 \
--exclude-unobserved-genotypes \
${LONG_READ_TUMOR_BAM} ${LONG_READ_NORMAL_BAM}
~~~

~~~
# Attempt to rescue variants uniquely present
# in the long read callset into “two-tech” calls
# by looking for them in short read alignments
/usr/bin/time --verbose freebayes \
--fasta-reference GRCh38_full_analysis_set_plus_decoy_hla.fa \
--variant-input ${LONG_READ_CALLSET_UNIQUE_VCF} \
--targets ${LONG_READ_CALLSET_UNIQUE_REGIONS_BED} \
--only-use-input-alleles --hwe-priors-off \
--binomial-obs-priors-off --allele-balance-priors-off \
--no-population-priors --legacy-gls \
--pooled-continuous --pooled-discrete \
--min-alternate-fraction 0 --min-alternate-count 1 \
--exclude-unobserved-genotypes \
${SHORT_READ_TUMOR_BAM} ${SHORT_READ_NORMAL_BAM}
~~~

The variants that were “rescued” using Freebayes are then filtered for sufficient evidence using the following bcftools (v1.20) expression, before they are added into the “two-tech” callset.

~~~
# at least 20x depth in both tumor and normal
# at least 2 or more reads supporting ALT allele in tumor
# no reads supporting ALT allele in normal
--include ‘FMT/DP[0]>=20 && FMT/DP[1]>=20 && SUM(FMT/AO[0:*])>=2 && SUM(FMT/AO[1:*])==0’
~~~

The entire workflow for “two-tech” truth set generation starting from the intersection of the long read and short read variant call sets is shown in Supplementary Figure 3.

The IGV screenshots (Supplementary Figures 8 to 23) containing both short and long read alignments from COLO829 tumor & normal samples show various examples of different classes of variants from the “two-tech” truth set generation process –

- Common (variants that are common between the short and long read call sets)
- LR_Origin (variants only in the long read call set validated by short read)
- ILMN_Origin (variants only in the short read call set validated by long read)
- Dropped (variants dropped from previously published high confidence truth set)

### 3. Variant Calling Performance Evaluation

The full command lines and exact tool versions that were used to run each of the variant callers on a single chromosome for performance evaluation are as follows –

#### Lancet1 v1.1.0_b6c9067

~~~
set -euxo pipefail
/usr/bin/time --verbose lancet \
      --num-threads “∼{numCpus}” \
      --normal “∼{normalBamCram}” \
      --tumor “∼{tumorBamCram}” \
      --ref “∼{referenceFasta}” \
      --reg “∼{contigRegion}” \
      --max-avg-cov 1000 \
      > tmp.lancet1_out.vcf 2> “stderr.∼{outputPrefix}.log” \ && bcftools reheader \
      --fai “∼{referenceFaidx}” “tmp.lancet1_out.vcf” \
| bcftools view -Oz -o “∼{outputPrefix}.vcf.gz” /dev/stdin \ && bcftools index --tbi “∼{outputPrefix}.vcf.gz”
~~~

#### Lancet2 v2.8.5-main-affb044c

~~~
set -euxo pipefail
# Increase max open files limit
ulimit -n 16384; ulimit -a
/usr/bin/time -v Lancet2 pipeline \
      --normal “∼{normalBamCram}” \
      --tumor “∼{tumorBamCram}” \
      --num-threads “∼{numCpus}” \
      --reference “∼{referenceFasta}” \
      --region “∼{contigRegion}” \
      --out-vcfgz “∼{outputPrefix}.vcf.gz” \
      |& tee “stderr.∼{outputPrefix}.log”
~~~

#### DeepSomatic v17_rc0_07012024

~~~
set -euxo pipefail
/usr/bin/time --verbose run_deepsomatic \
      --model_type=WGS --ref=“∼{referenceFasta}” \
      --reads_normal=“∼{normalBamCram}” \
      --reads_tumor=“∼{tumorBamCram}” \
      --sample_name_tumor=∼{tumorName} \
      --sample_name_normal=∼{normalName} \
      --num_shards=“∼{numCpus}” \
      --regions=“∼{contigRegion}” \
      --output_vcf=“∼{outputPrefix}.vcf.gz” \
      |& tee “stderr.∼{outputPrefix}.log”
~~~

#### Mutect2 gatk-v4.6.0.0

~~~
set -euxo pipefail
# Mutect2 needs ref dict and exact sample name in RG header gatk CreateSequenceDictionary --REFERENCE “∼{referenceFasta}”
gatk GetSampleName -I “∼{normalBamCram}” -O normalName.txt && \ gatk GetSampleName -I “∼{tumorBamCram}” -O tumorName.txt
JAVA_OPTS=“-Xmx∼{memoryInGb}g -XX:ParallelGCThreads=∼{numCpus}”
/usr/bin/time --verbose gatk Mutect2 \
      --java-options “∼{JAVA_OPTS}” \
      --native-pair-hmm-threads “∼{numCpus}” \
      --input “∼{normalBamCram}” \
      --input “∼{tumorBamCram}” \
      --reference “∼{referenceFasta}” \
      --intervals “∼{contigRegion}” \
      --normal-sample “$(cat normalName.txt)” \
      --tumor-sample “$(cat tumorName.txt)” \
      --output “∼{outputPrefix}.raw.vcf.gz” \
      --annotation-group StandardMutectAnnotation \
      |& tee “stderr.∼{outputPrefix}.log”
/usr/bin/time --verbose gatk FilterMutectCalls \
      --java-options “∼{JAVA_OPTS}” \
      --reference “∼{referenceFasta}” \
      --variant “∼{outputPrefix}.raw.vcf.gz” \
      --output “∼{outputPrefix}.vcf.gz” \
      |& tee “stderr.∼{outputPrefix}.log”
~~~

#### Strelka2 v2.9.10-manta-v1.6.0

~~~
set -euxo pipefail
mkdir -p “∼{outputPrefix}.manta_runDir” \
         “∼{outputPrefix}.strelka_runDir”
chromRegex=“^∼{contigRegion}\t”
printf “%s\t0\t%s\n” \
       “∼{contigRegion}” \
       “$(grep -P ${chromRegex}
    ∼{referenceFaidx} | cut -f2)” \
| bgzip -c >| call_regions.bed.gz \
&& tabix -p bed call_regions.bed.gz
configManta.py \
      --referenceFasta “∼{referenceFasta}” \
      --normalBam “∼{normalBamCram}” \
      --tumourBam “∼{tumorBamCram}” \
      --runDir “manta_runDir” \
      --callRegions “call_regions.bed.gz” \
&& /usr/bin/time --verbose \
      “manta_runDir/runWorkflow.py” \
      --mode local --jobs “∼{numCpus}” \
      |& tee “stderr.∼{outputPrefix}.log”
configureStrelkaSomaticWorkflow.py \
      --referenceFasta “∼{referenceFasta}” \
      --normalBam “∼{normalBamCram}” \
      --tumourBam “∼{tumorBamCram}” \
      --runDir “strelka_runDir” \
      --callRegions “call_regions.bed.gz” \
      --indelCandidates \
“manta_runDir/results/variants/candidateSmallIndels.vcf.gz” \
&& /usr/bin/time --verbose \
      “strelka_runDir/runWorkflow.py” \
      --mode local --jobs “∼{numCpus}” \
      |& tee “stderr.∼{outputPrefix}.log”
~~~

#### Varnet v1.1.0

~~~
set -euxo pipefail
mkdir -p “∼{outputPrefix}”
chromRegex=“^∼{contigRegion}\t”
printf “%s\t0\t%s\n” \
       “∼{contigRegion}” \
       “$(grep -P ${chromRegex} ∼{referenceFaidx} | cut -f2)” \
       >| region.bed
/usr/bin/time --verbose python /VarNet/filter.py \
       --processes “∼{numCpus}” \
       --sample_name “∼{tumorName}” \
       --reference “∼{referenceFasta}” \
       --region_bed “region.bed” \
       --normal_bam “∼{normalBamCram}” \
       --tumor_bam “∼{tumorBamCram}” \
       --output_dir “∼{outputPrefix}” \
       |& tee “stderr.∼{outputPrefix}.log” \
&& /usr/bin/time --verbose python /VarNet/predict.py \
       --processes “∼{numCpus}” \
       --sample_name “∼{tumorName}” \
       --reference “∼{referenceFasta}” \
       --normal_bam “∼{normalBamCram}” \
       --tumor_bam “∼{tumorBamCram}” \
       --output_dir “∼{outputPrefix}” \
       |& tee “stderr.∼{outputPrefix}.log”
~~~

The per chromosome variant calls from each caller was then normalized and combined using the following bcftools (v1.20) commands

~~~
# Normalize
bcftools reheader --fai “∼{referenceFaidx}” ∼{rawVcfGz} \
| bcftools norm -Oz -o “∼{outputVcf}” \
           --fasta-ref “∼{referenceFasta}” --check-ref s \ && bcftools index --tbi “∼{outputVcf}”
# Combine
bcftools concat -Ou ∼{sep=’’ vcfs} \
| bcftools sort -m12G -Oz -o “∼{outputPrefix}.vcf.gz” \
&& bcftools index --tbi “∼{outputPrefix}.vcf.gz”
~~~

Java v1.9 and RTG vcfeval v3.12.1 were then used to benchmark each variant caller against the truth set using the following command –

~~~
rtg vcfeval \
    --template GRCh38_full_analysis_set_plus_decoy_hla.sdf \
    --baseline ${TRUTH_VCF} \
    --calls ${CALLSET_VCF} \
    --output ${OUTPUT_DIRECTORY} \
    --all-records --sample=ALT,ALT \
    --vcf-score-field=QUAL \
    --output-mode=annotate
~~~

The full workflow used to perform variant caller performance evaluation which also contains additional scripts to postprocess tool VCFs and generate an interactive benchmarking HTML report is available at the following git repository – https://bitbucket.nygenome.org/projects/COMPBIO-INTERNAL/repos/lancet2_manusc ript/browse.

For convenience the interactive benchmarking reports generated from the analysis are available at the following links –

Using the Two-Tech Truthset

https://storage.googleapis.com/lancet2-paper/reports/vs_two_tech_truth/HCC1187/H CC1187_WholeGenome.html

https://storage.googleapis.com/lancet2-paper/reports/vs_two_tech_truth/HCC1143/H CC1143_WholeGenome.html

https://storage.googleapis.com/lancet2-paper/reports/vs_two_tech_truth/COLO829/C OLO829_WholeGenome.html

https://storage.googleapis.com/lancet2-paper/reports/vs_two_tech_truth/HCC1395/H CC1395_WholeGenome.html

Using the previously published Illumina only High Confidence Truthset

https://storage.googleapis.com/lancet2-paper/reports/vs_ilmn_hc_truth/HCC1187/HC C1187_WholeGenome.html

https://storage.googleapis.com/lancet2-paper/reports/vs_ilmn_hc_truth/HCC1143/HC C1143_WholeGenome.html

https://storage.googleapis.com/lancet2-paper/reports/vs_ilmn_hc_truth/COLO829/CO LO829_WholeGenome.html

https://storage.googleapis.com/lancet2-paper/reports/vs_ilmn_hc_truth/HCC1395/HC C1395_WholeGenome.html

### 4. Runtime Performance Evaluation

The same command lines were used to perform runtime performance evaluation for all the variant callers as shown previously in supplementary section 3. Two different benchmarking experiments were performed on Google Cloud using HCC1395 tumor and matched normal (HCC1395BL) Illumina datasets with the following variables – 1.) constant coverage dataset with increasing CPU core counts. 2.) constant CPU core count with increasing dataset coverages.

To study the scalability of variant callers, we performed the first benchmarking experiment on n2-highmem machines with varying CPU core count (2, 4, 8, 16, 32, 64, 128 CPU cores were used) to process chromosome 20 of HCC1395 tumor and matched normal data, both downsampled to 64x coverage. To study the impact of sample coverage, the second benchmarking experiment was performed on n2-highmem-128 machines with increasing coverage (64x/64x, 128x/128x, 256x/256x, 512x/512x, 1024x/1024x tumor/normal coverages were used) to process chromosome 20 of HCC1395 tumor and matched normal data.

### 5. Graph Visualization of Variants using Sequence Tube Map

The following steps describe the workflow to visualize Lancet2 variants of interest in graph space using the Sequence Tube Map framework.

1. Install prerequisite tools – Lancet2 (https://github.com/nygenome/Lancet2), samtools, bcftools, vg version 1.59.0, and jq, and ensure that they are available as commands that can be executed in the environment PATH.
2. Install Sequence Tube Map (https://github.com/vgteam/sequenceTubeMap) version `0452ecb82d057372e359a9b456d789336e5ab8a1`.
3. Use the Lancet2’s prep_stm_viz.sh script to run Lancet2 on a small set of variants of interest that need to be visualized in Sequence Tube Map. The script will run Lancet2 using the --graph-dir flag to generate GFA formatted sequence graphs for each variant of interest. Local VG graphs and indices required to load the sample reads along with the Lancet2 graph are then constructed, enabling simplified use with the “custom” data option in the Sequence Tube Map interface.
4. After running the Sequence Tube Map Server as detailed in the Tube Map Readme, set “Data” to “custom” and “BED file” to “index.bed”. Pick a “Region”(or variant) of interest and hit “Go” to visualize.

Sequence Tube Map demo server containing somatic variants from COLO829 tumor/normal pair sample – https://shorturl.at/JQZsy

## Supplementary Figures

**Supplementary Figure 1:**
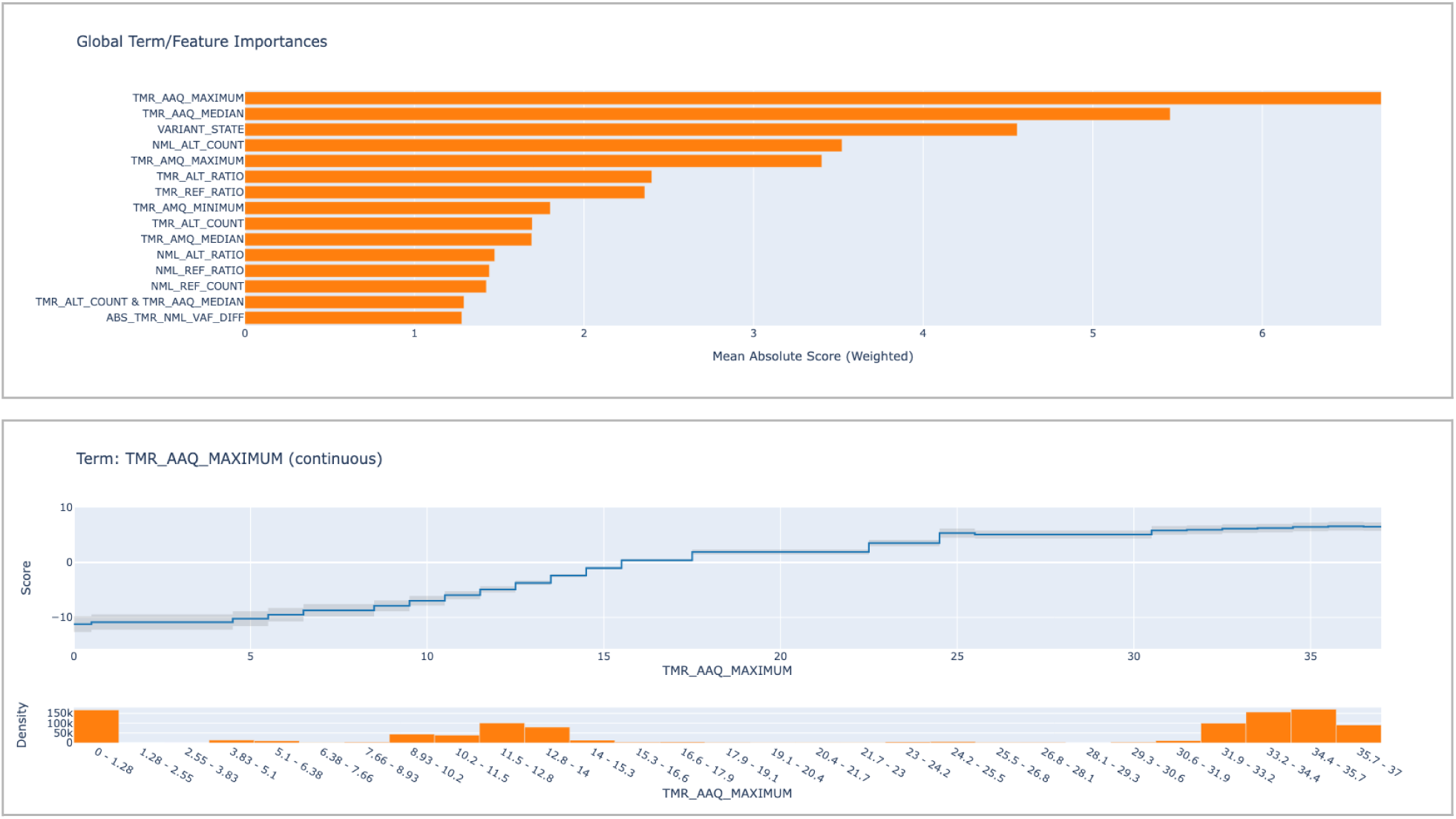
Global Explanation to the Somatic Glassbox Model. The first panel shows the relative term/feature importances and their mean weighted scores learned by the Somatic Glassbox model. The second panel shows the expected feature response curve for a single feature in the model (TME_AAQ_MAXIMUM). The X axis shows the values of the feature in question with the density histogram showing its distribution in the training set. The Y axis represents the log probability score learned by the model at a particular X-axis value.

**Supplementary Figure 2:**
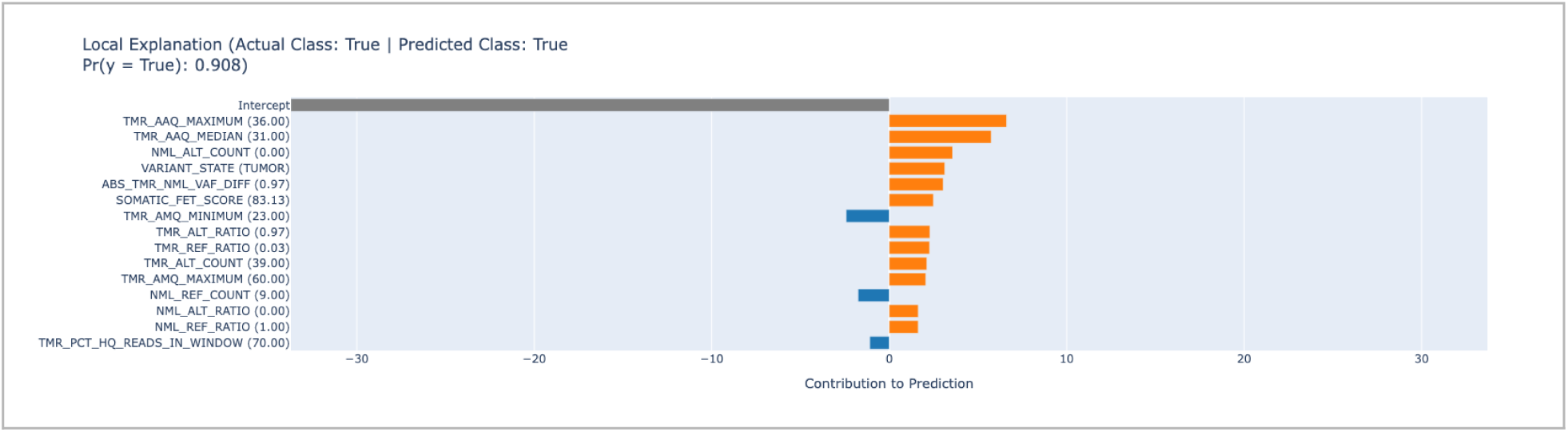
Local Explanation to the Somatic Glassbox Model. Shows the individual contribution of the most important features used by the model to arrive at its final prediction and probability value for the classification given a single variant. The feature values of the variant in question are shown in brackets along the Y-axis.

**Supplementary Figure 3:**
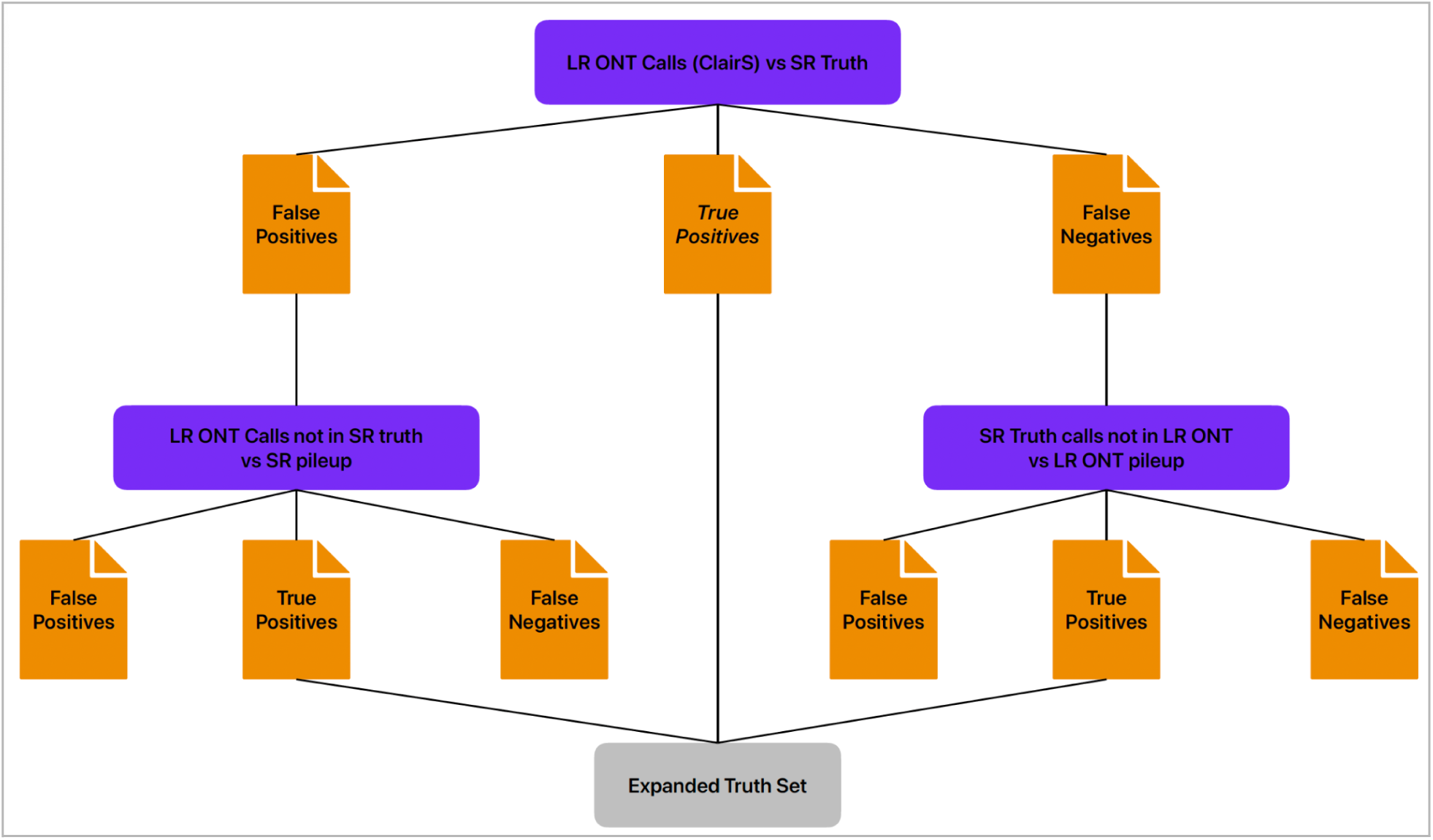
Flowchart showing a high level overview of the process used to “rescue” variants unique to a single technology into the “two-tech” truthset. All intersections between two different call sets were performed using RTG vcfeval v3.12.1. The terms “true positives”, “false positives” and “false negatives” refer to the output VCFs generated by vcfeval in the split output mode.

**Supplementary Figure 4:**
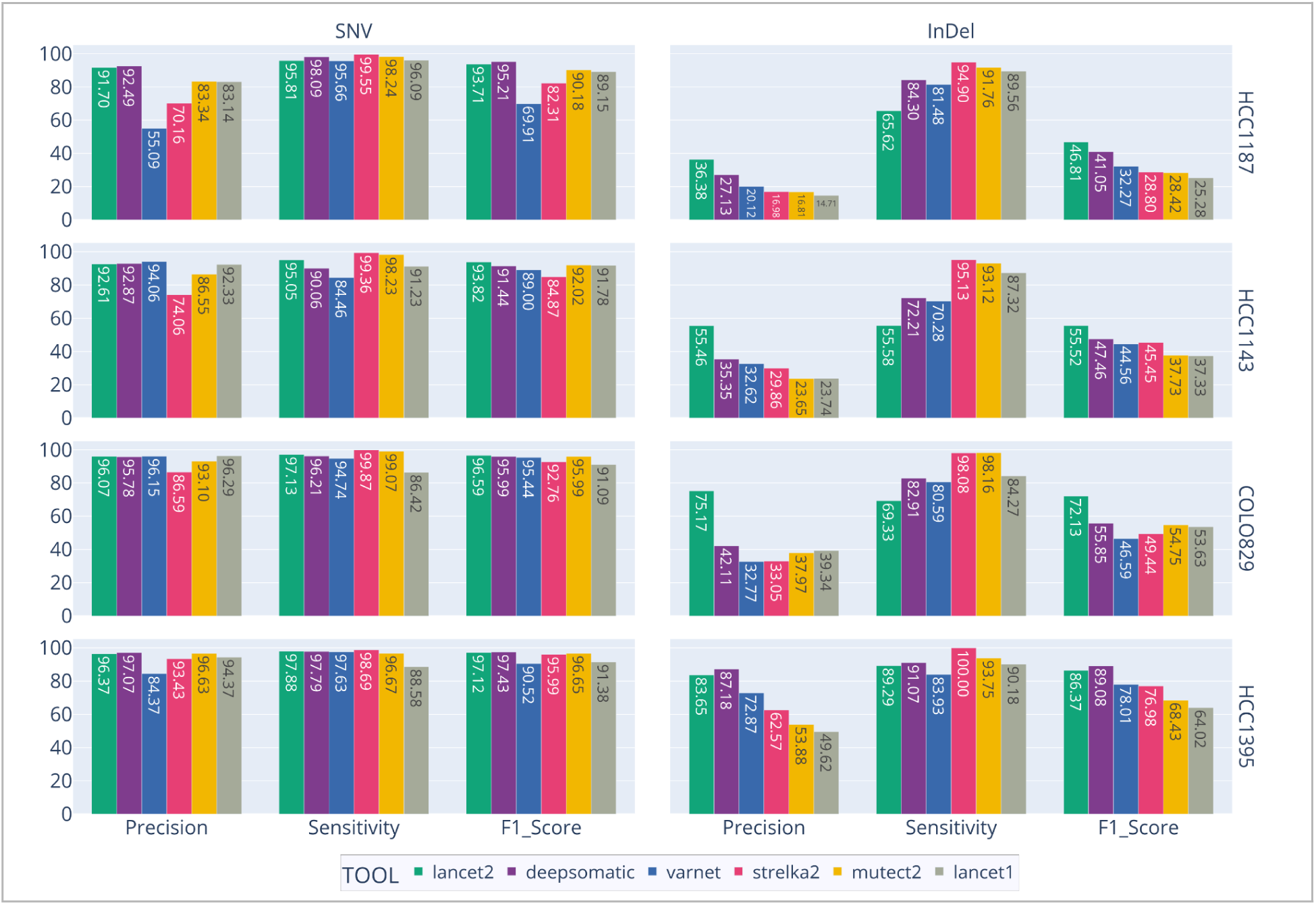
Comparison of variant calling performance metrics (precision, sensitivity, and F1 score) for multiple tools (Lancet2, DeepSomatic, VarNet, Strelka2, Mutect2, Lancet1) benchmarked against previously published high-confidence truth sets. Each panel represents a distinct variant type (SNV, InDel) and each row corresponds to one of four cancer cell lines (HCC1187, HCC1143, COLO829, HCC1395). Bars indicate the performance metric values for each tool, with numerical labels displayed.

**Supplementary Figure 5:**
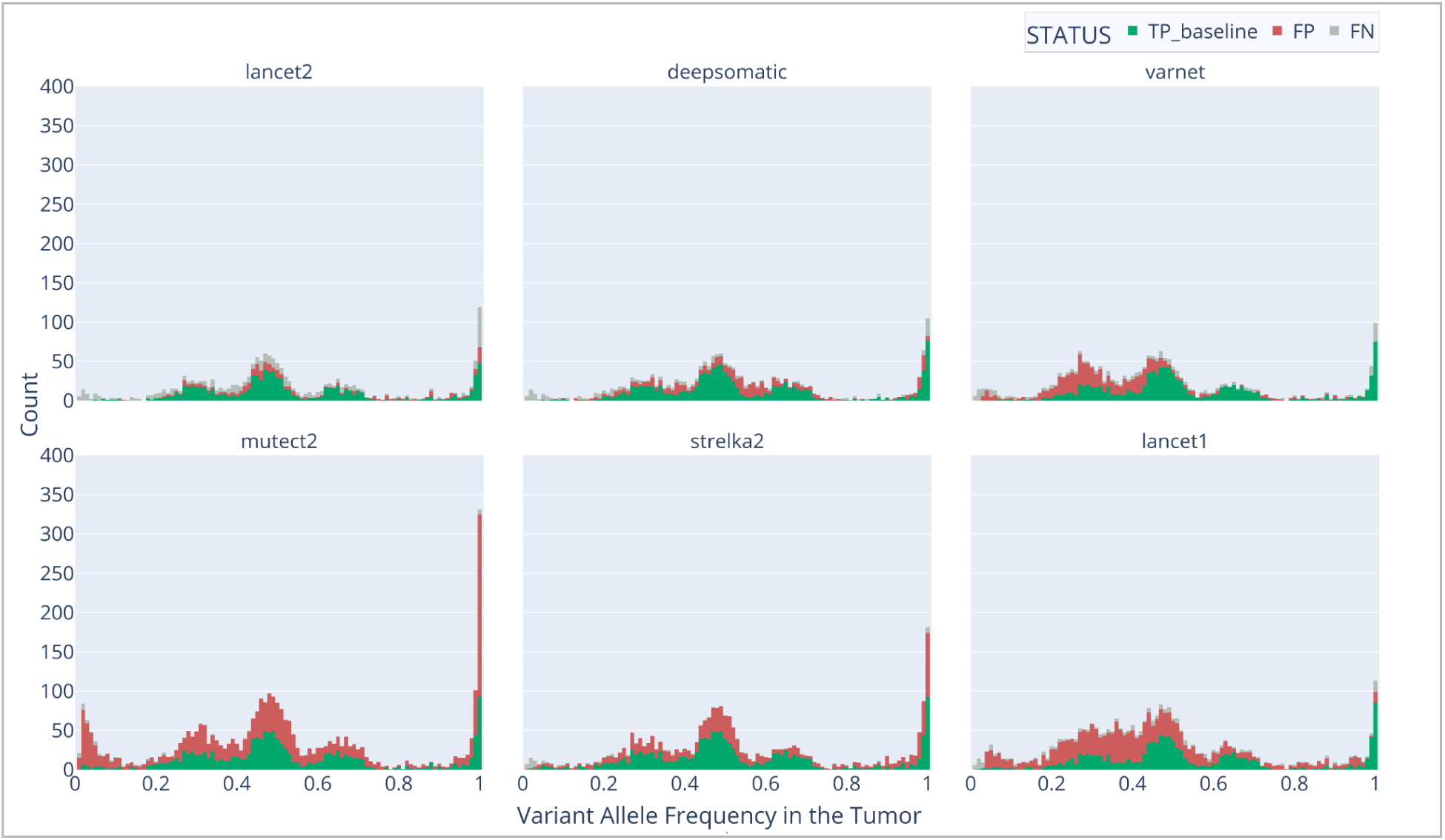
Variant allele frequency (VAF) distributions of true positive calls, false positives, and false negatives for InDel variants identified by multiple variant callers (Lancet2, DeepSomatic, VarNet, Strelka2, Mutect2, and Lancet1) in the COLO829 dataset. Each panel shows the count of variants at different VAF levels stratified by call status for a single calling tool. VAF is extracted from the truth set calls for false negative calls and extracted from the variant caller output for true positive and false positive calls.

**Supplementary Figure 6:**
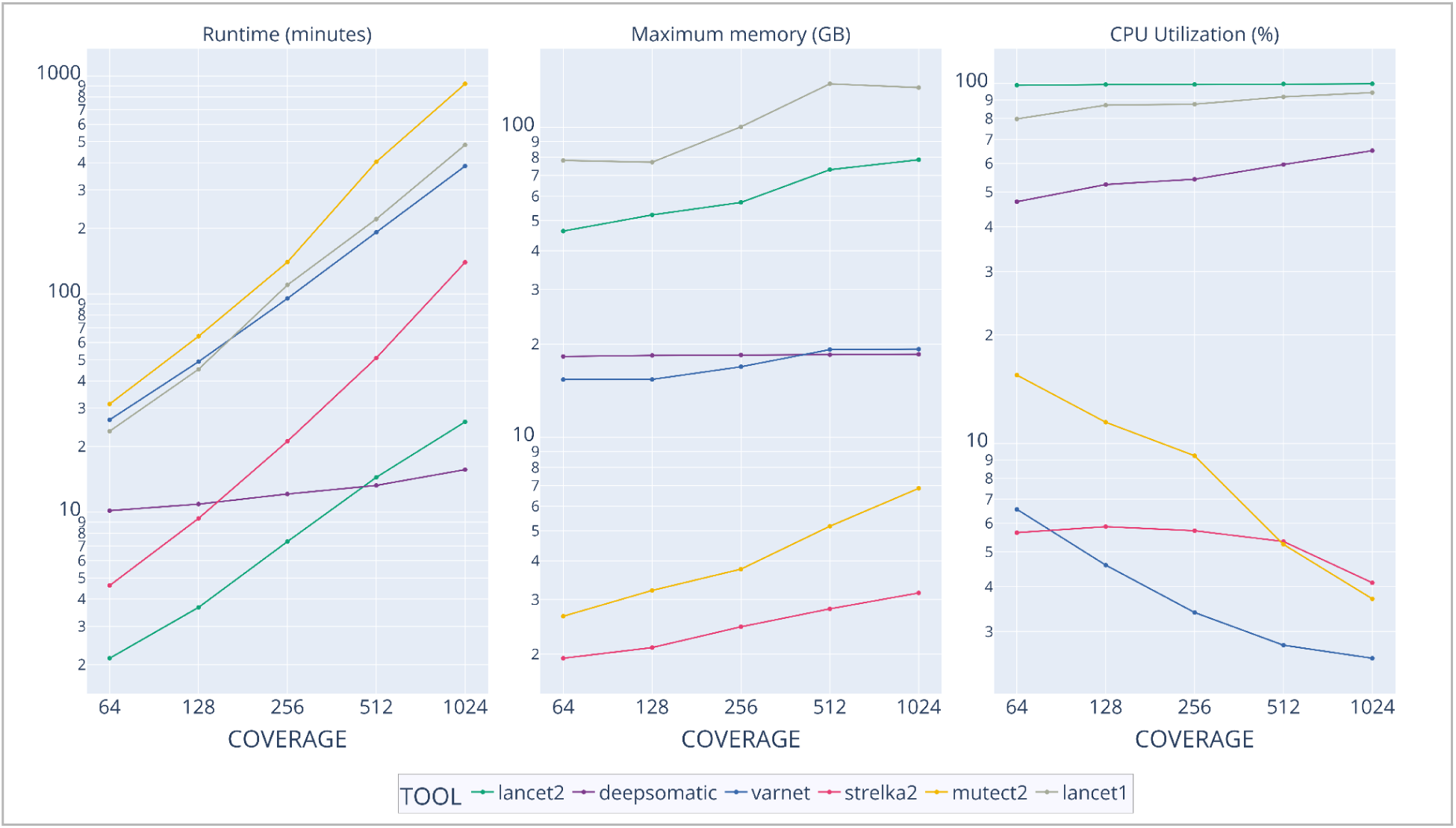
Runtime Performance by Coverage: Benchmarked on Google Cloud with a 128 core (n2-highmem-128) machine on chr20 HCC1395 vs HCC1395BL both at same coverage. Three line plots showing runtime in minutes, maximum memory usage in GB, and percent CPU utilization for multiple variant calling tools (Lancet2, Lancet1, DeepSomatic, Mutect2, Strelka2, VarNet) as sequencing coverage increases. The x-axes represent increasing coverage levels, and the y-axes represent the respective metric values. Each line corresponds to a different tool.

**Supplementary Figure 7:**
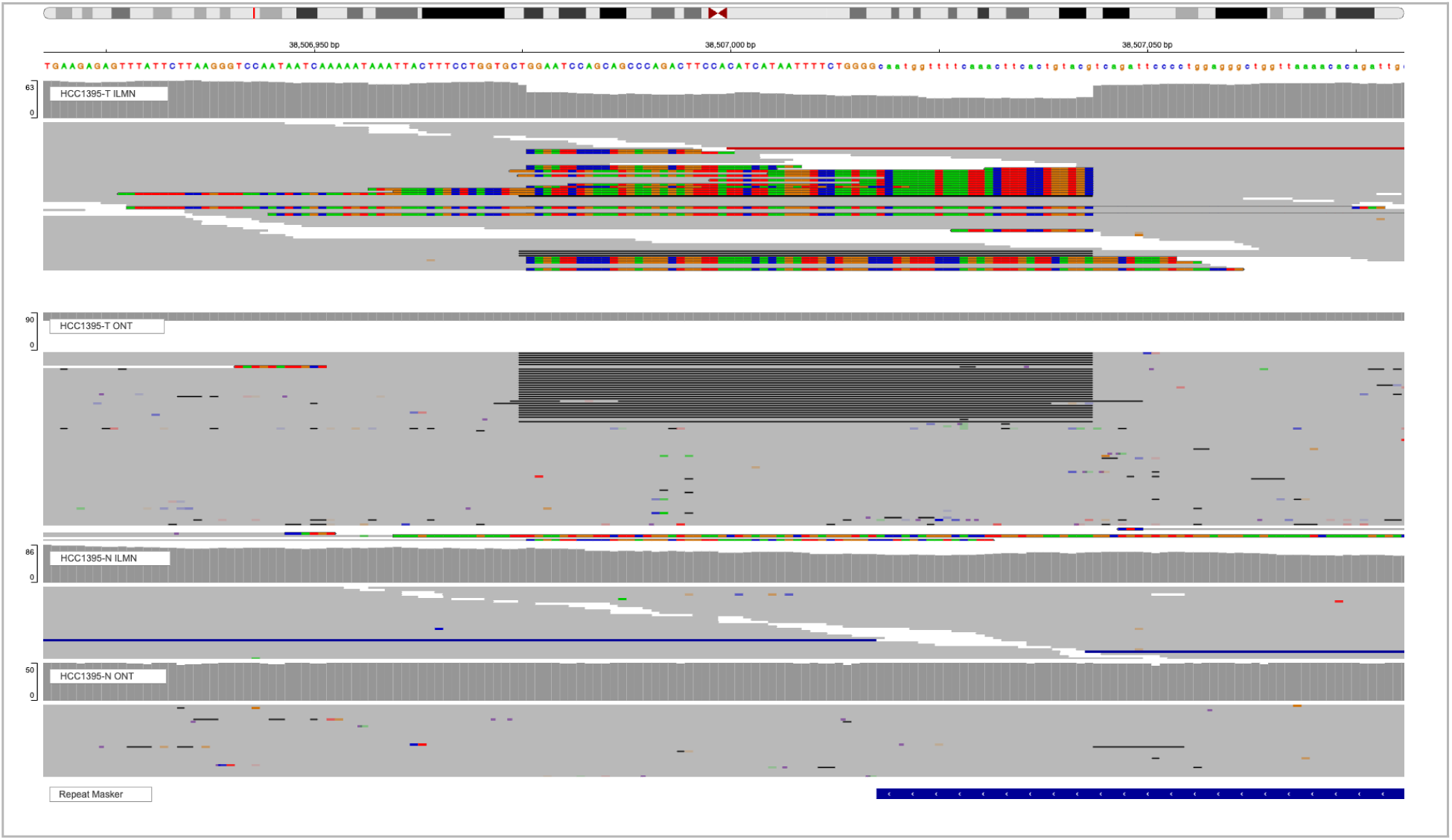
Integrated Genome Viewer (IGV) visualization of the same 70-bp somatic deletion in HCC1395 tumor and normal cell lines as in Figure 5 with alignment tracks for short read tumor, long read tumor, short read normal and long read normal samples respectively.

**Supplementary Figure 8:**
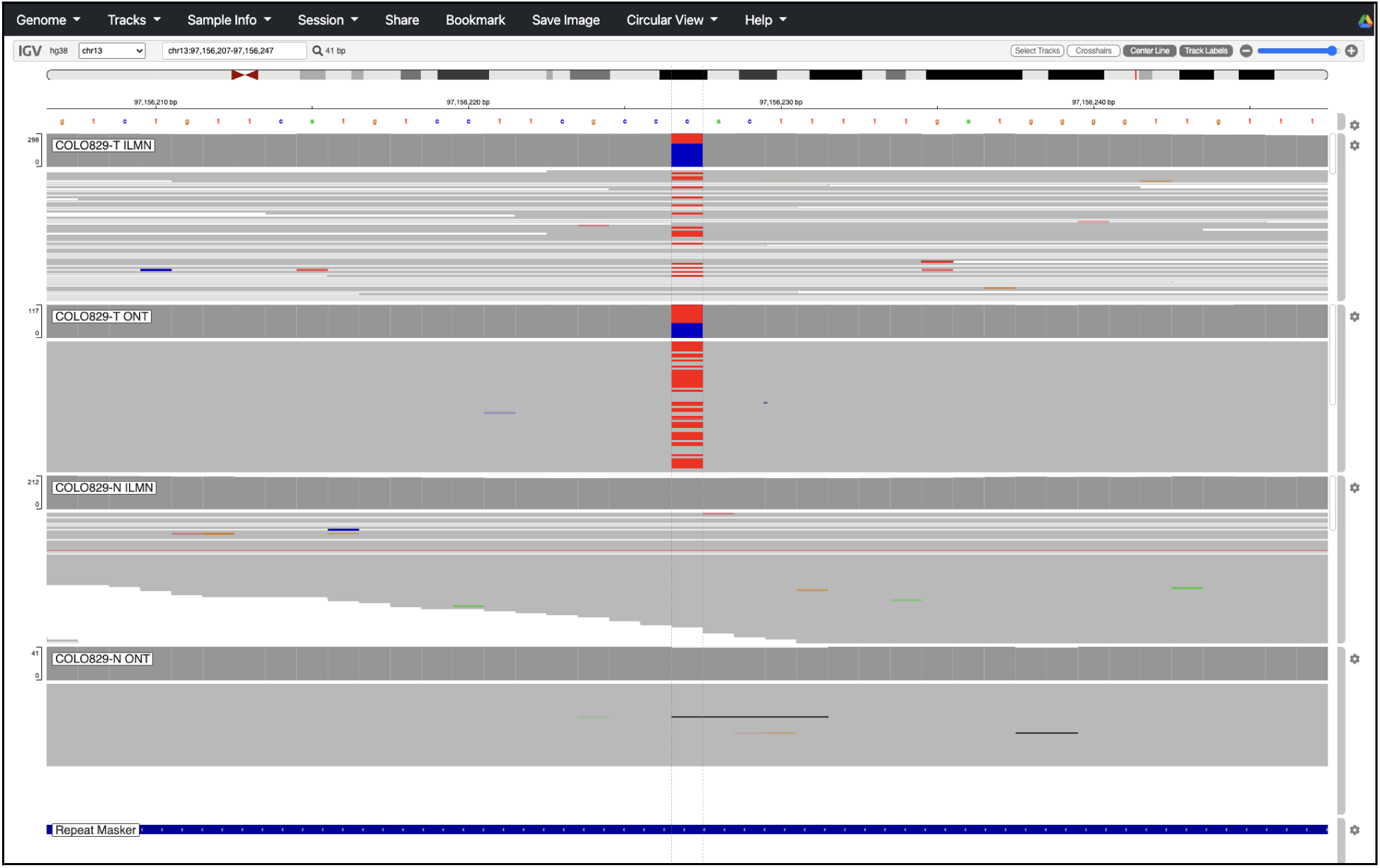
C → T somatic SNV in COLO829 at chr13:97156227 common between the short and long read call sets.

**Supplementary Figure 9:**
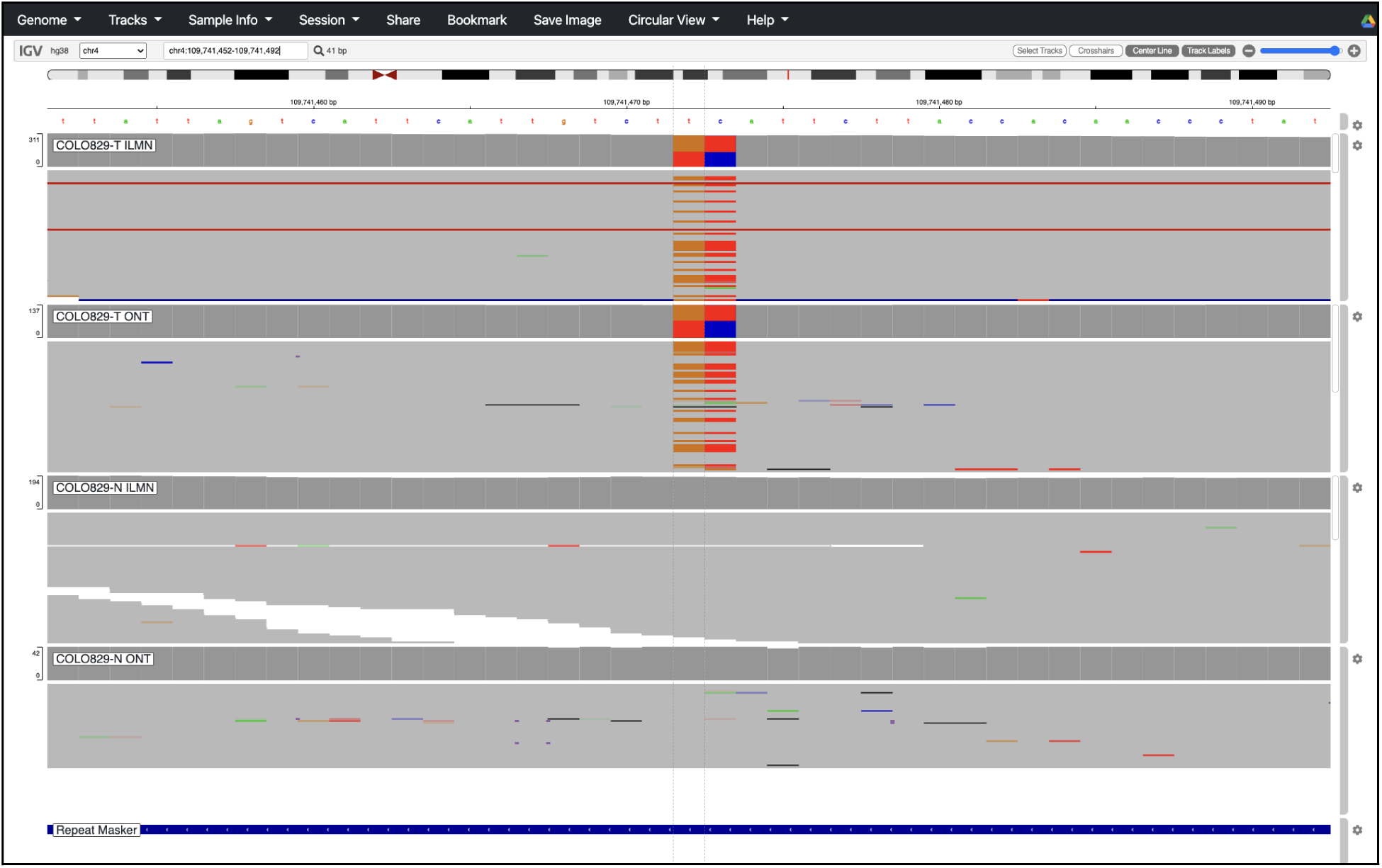
TC → GT somatic MNV in COLO829 at chr4:109741472 common between the short and long read call sets.

**Supplementary Figure 10:**
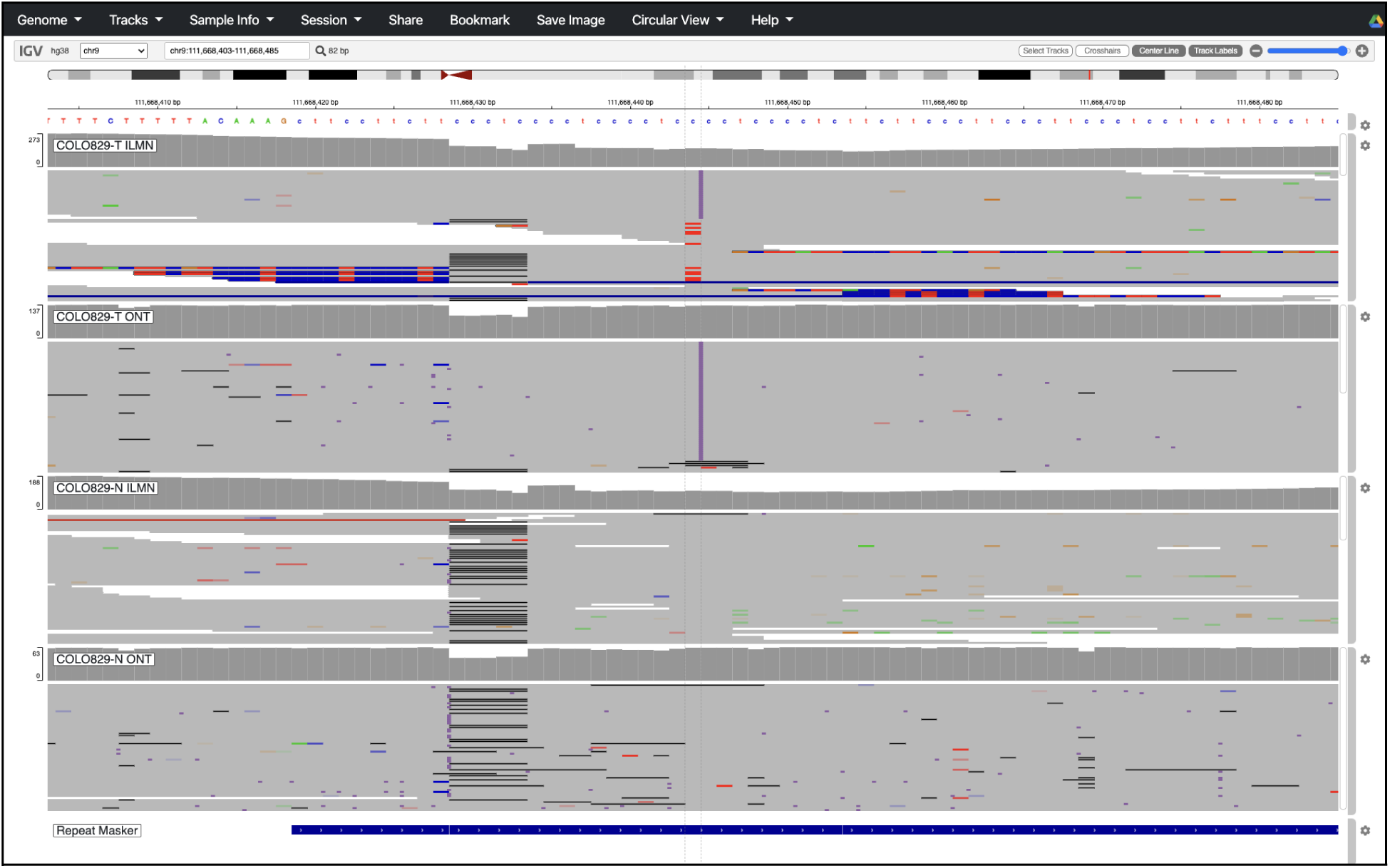
15 bp somatic INS in COLO829 at chr9:111668444 common between the short and long read call sets.

**Supplementary Figure 11:**
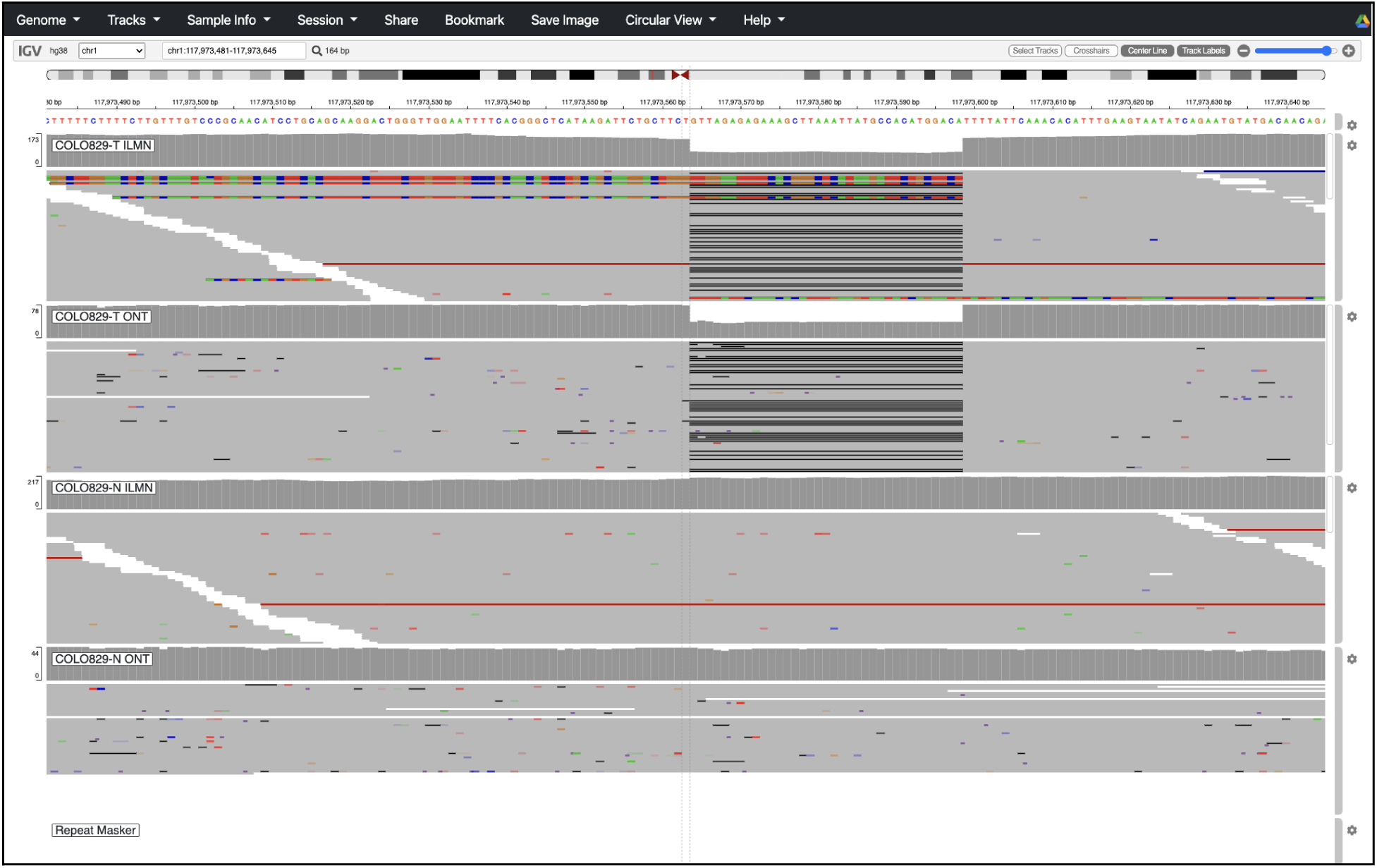
35 bp somatic DEL in COLO829 at chr1:117973563 common between the short and long read call sets.

**Supplementary Figure 12:**
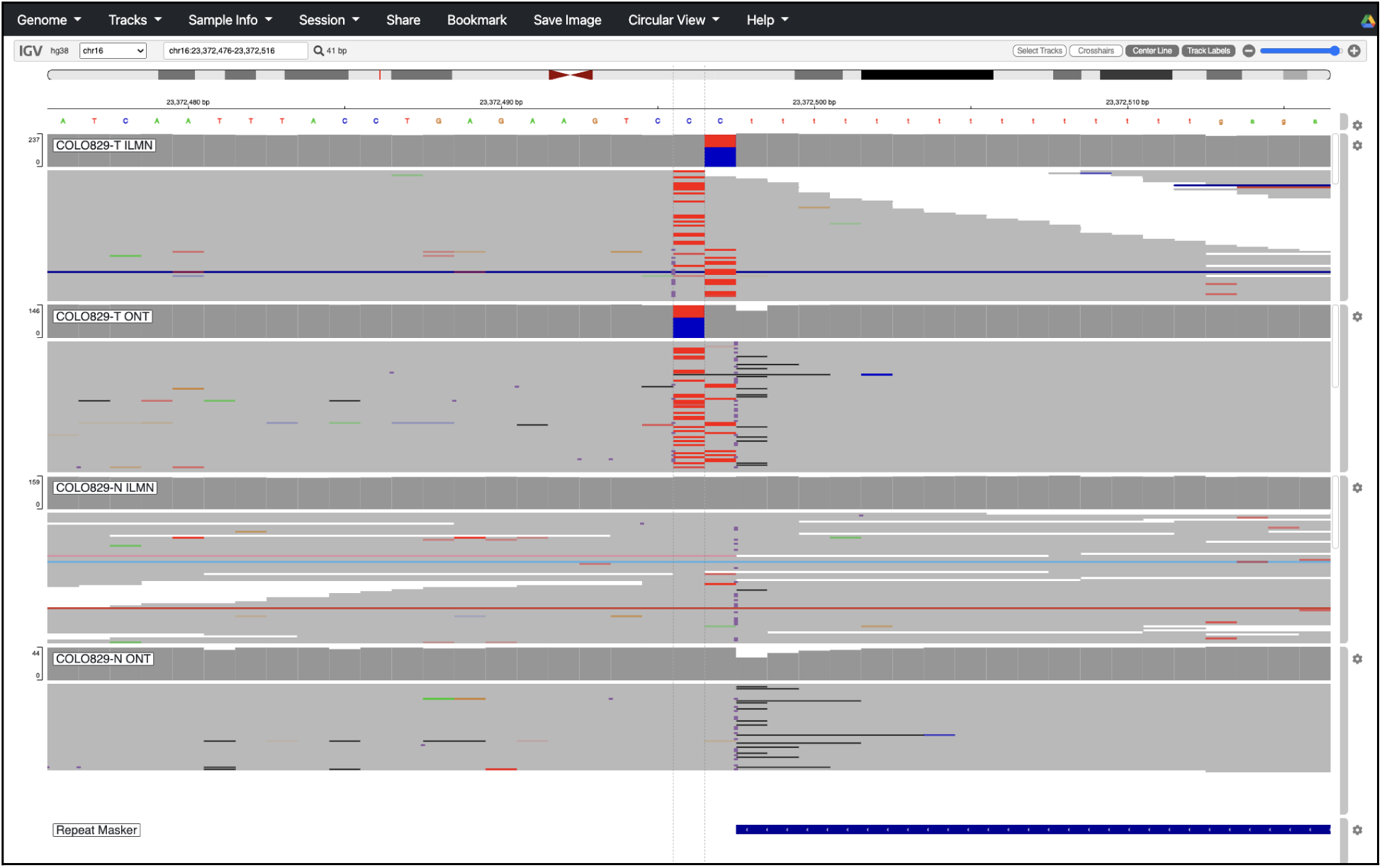
C → T somatic SNV in COLO829 at chr16:23372496 found in the long read call set and validated by short read alignments.

**Supplementary Figure 13:**
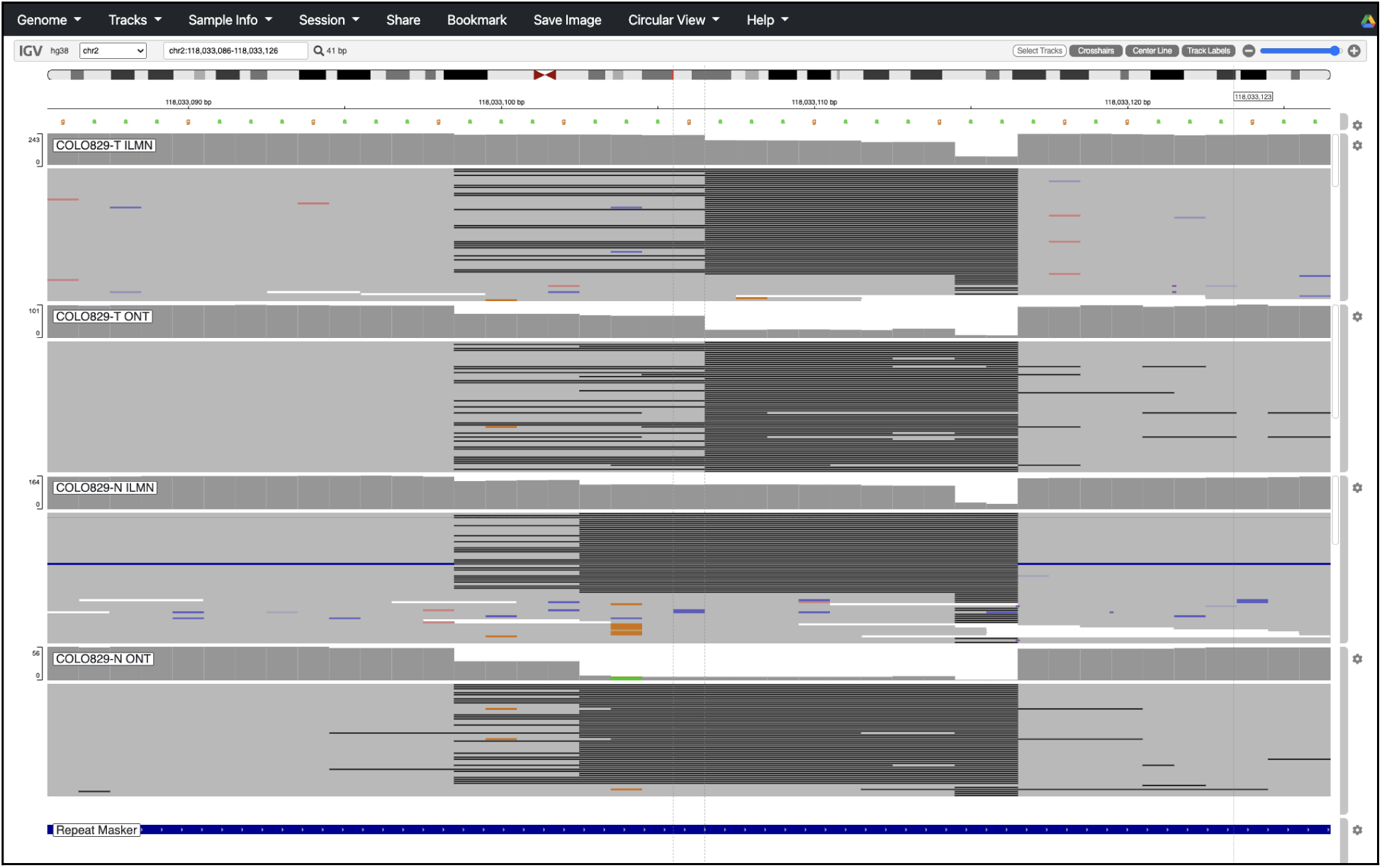
10 bp somatic DEL in COLO829 at chr2:118033106 found in the long read call set and validated by short read alignments.

**Supplementary Figure 14:**
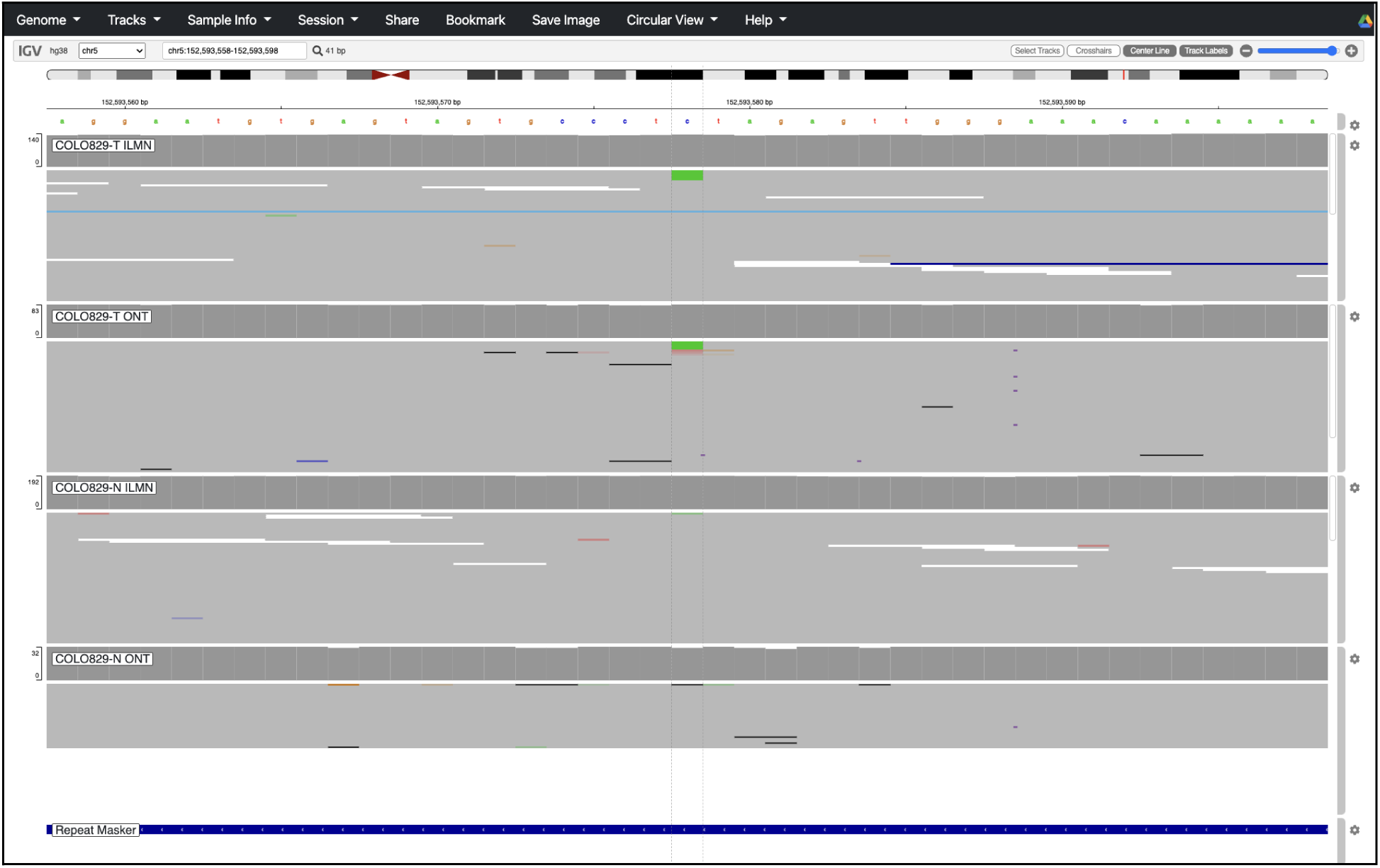
C → A somatic SNV in COLO829 at chr5:152593578 found in the short read call set and validated by long read alignments.

**Supplementary Figure 15:**
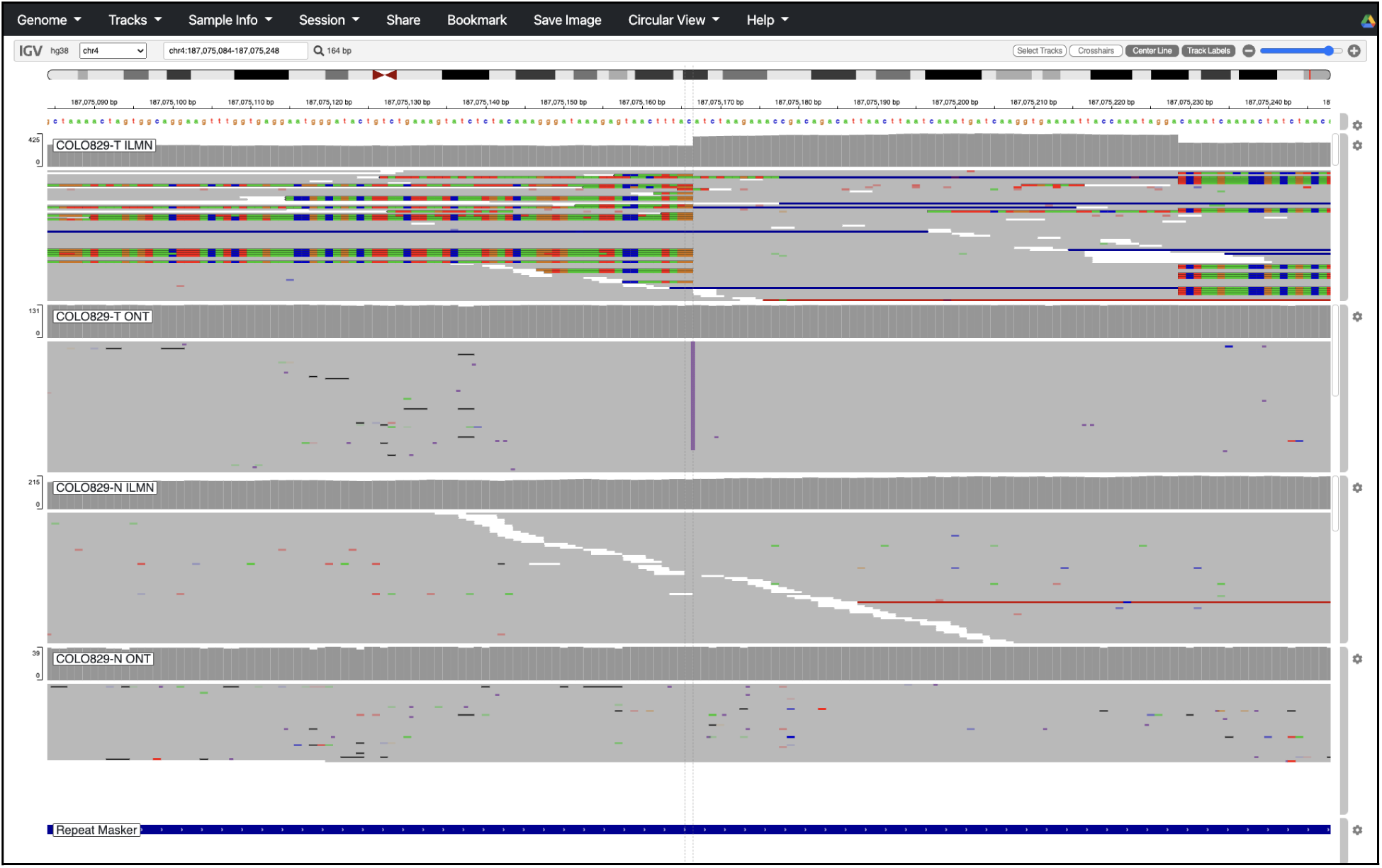
61 bp somatic INS in COLO829 at chr4:187075166 found in the short read call set and validated by long read alignments.

**Supplementary Figure 16:**
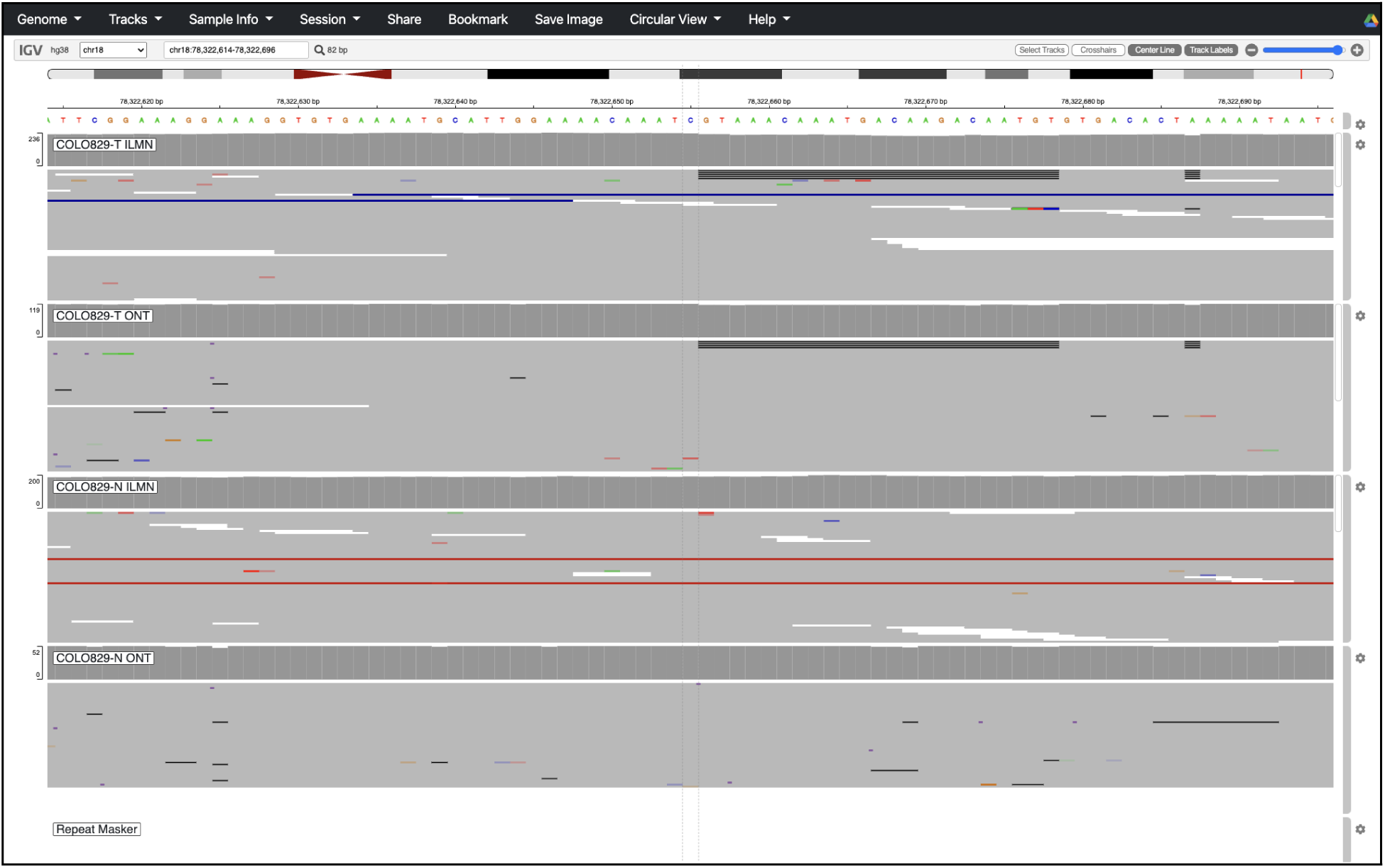
23 bp somatic DEL in COLO829 at chr18:78322655 found in the short read call set and validated by long read alignments.

**Supplementary Figure 17:**
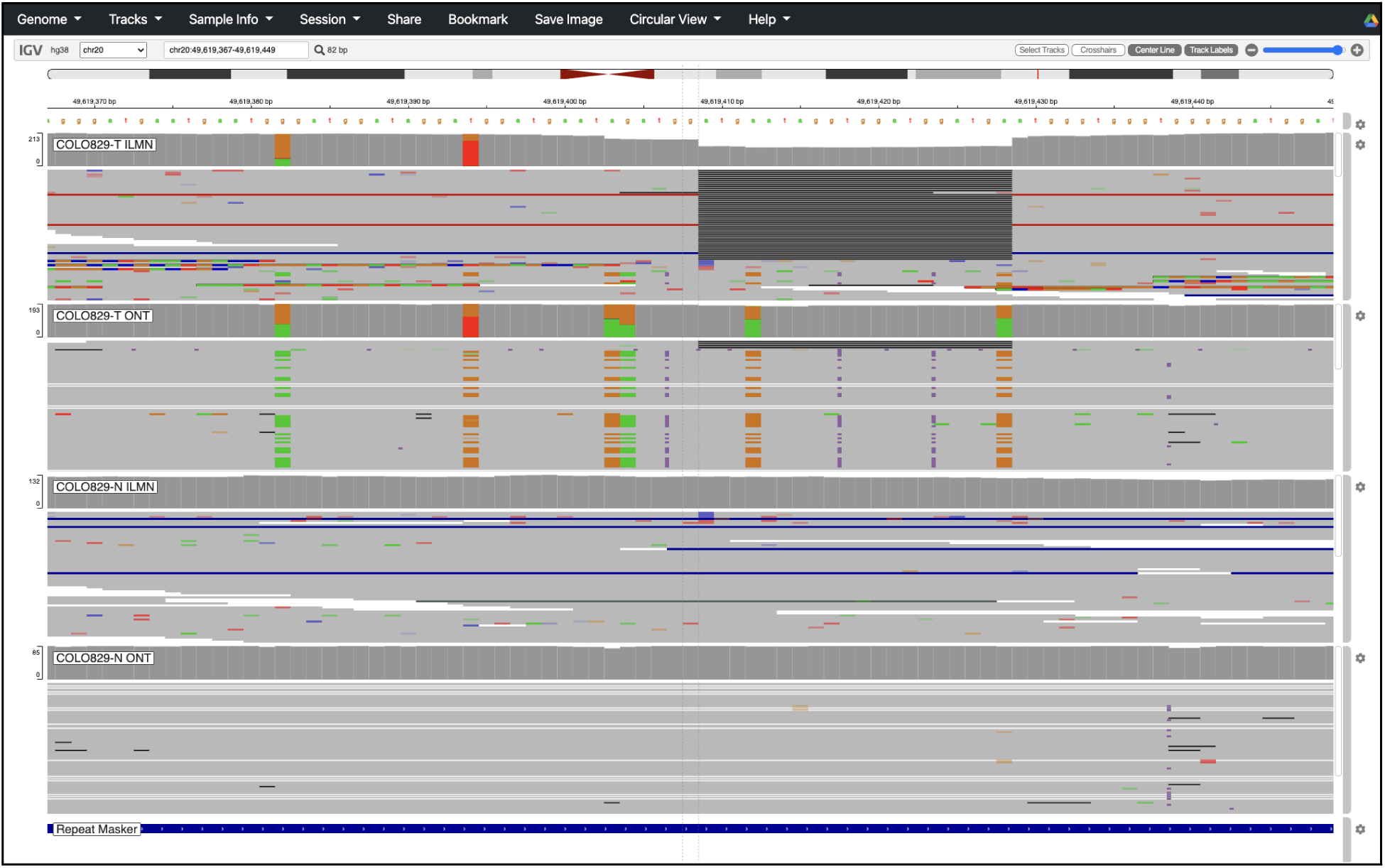
20 bp somatic DEL in COLO829 at chr20:49619408 found in the short read call set and validated by long read alignments.

**Supplementary Figure 18:**
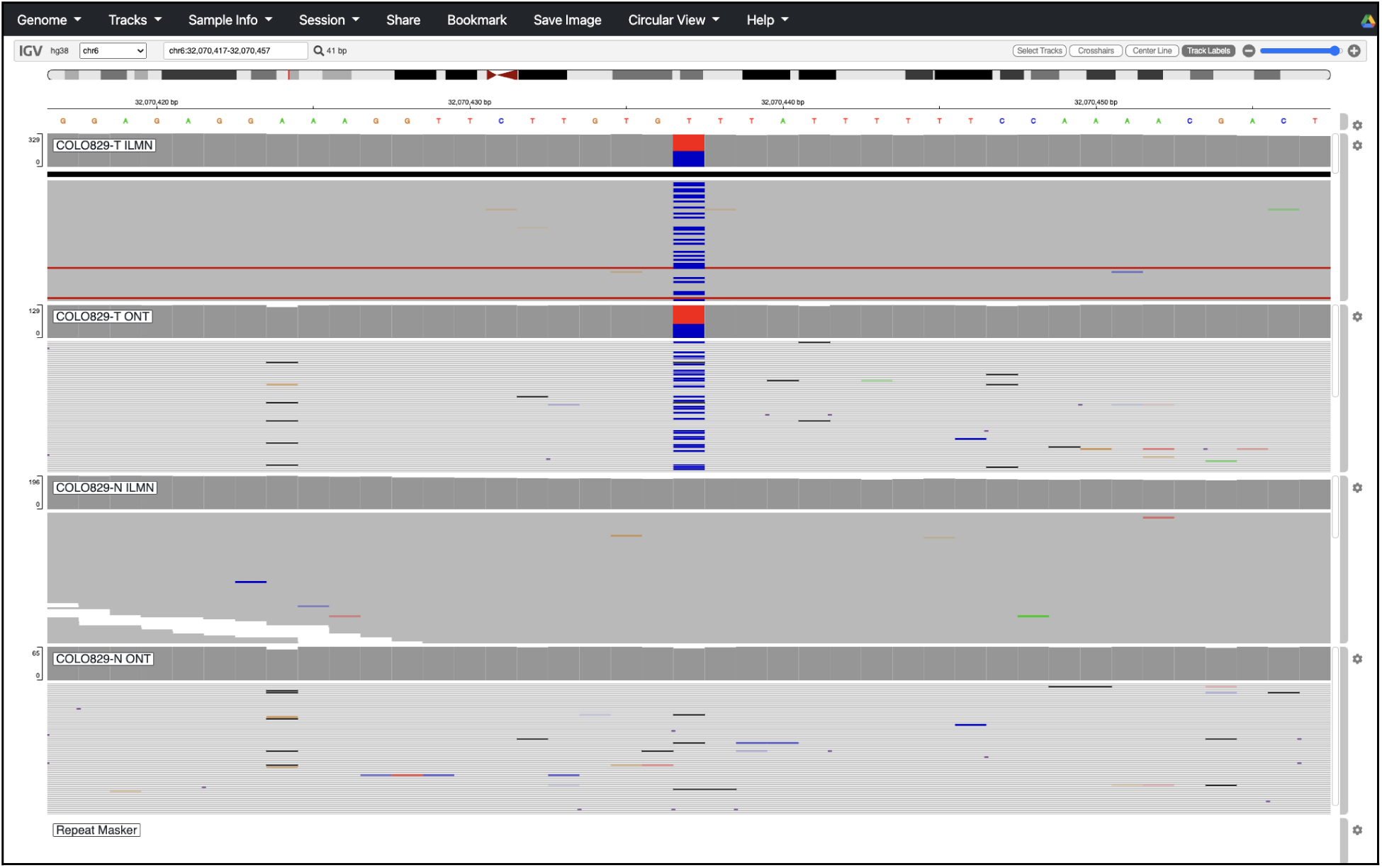
T → C putative somatic SNV in COLO829 at chr6:32070437 dropped from the previous high confidence short read truth set. Reason – only ambiguous mapping quality zero long read alignments support ALT allele.

**Supplementary Figure 19:**
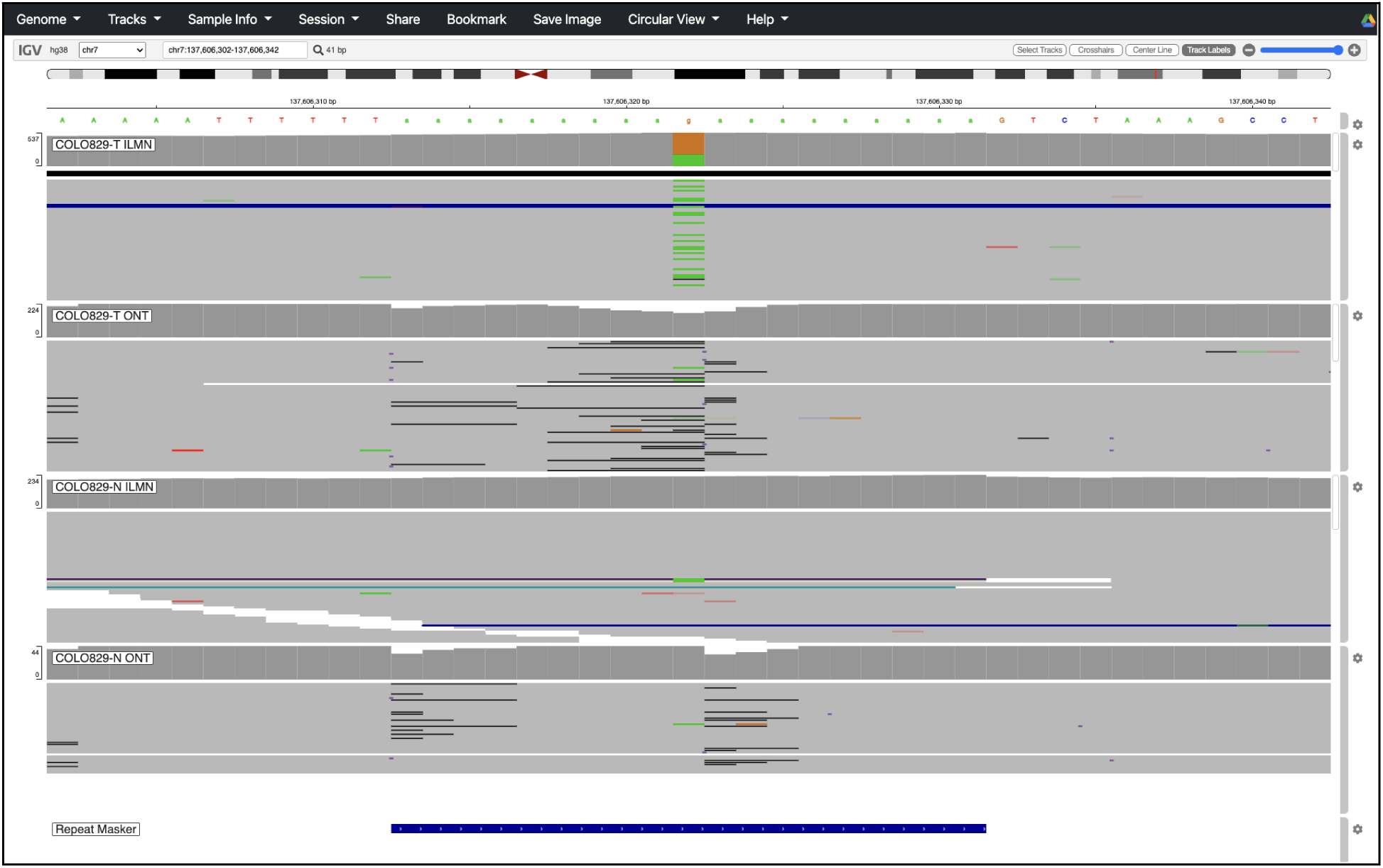
G → A putative somatic SNV in COLO829 at chr7:137606322 dropped from the previous high confidence short read truth set. Reason – ALT allele seen in normal sample.

**Supplementary Figure 20:**
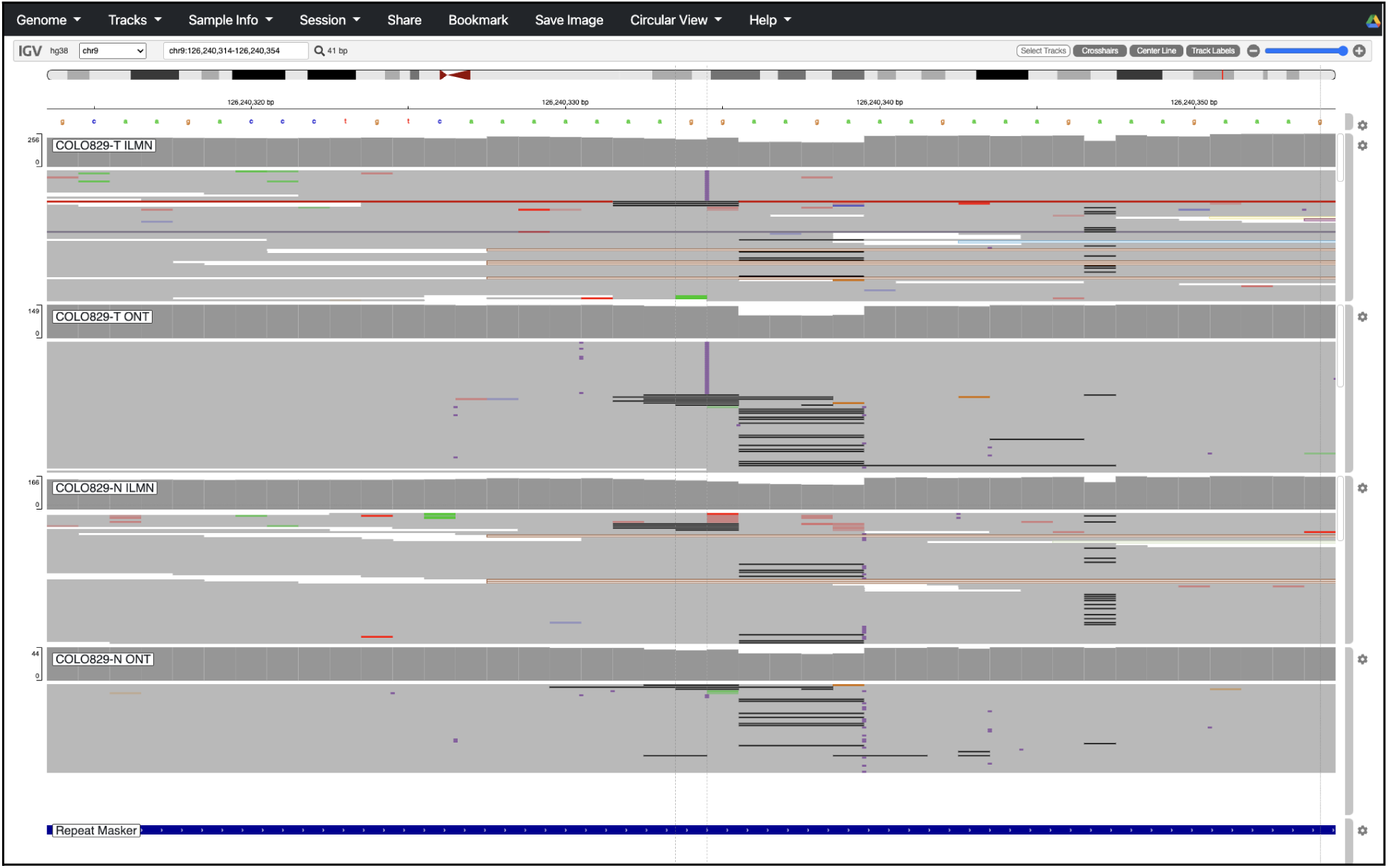
11 bp putative somatic INS in COLO829 at chr9:126240334 dropped from the previous high confidence short read truth set. Reason – ALT allele seen in normal sample.

**Supplementary Figure 21:**
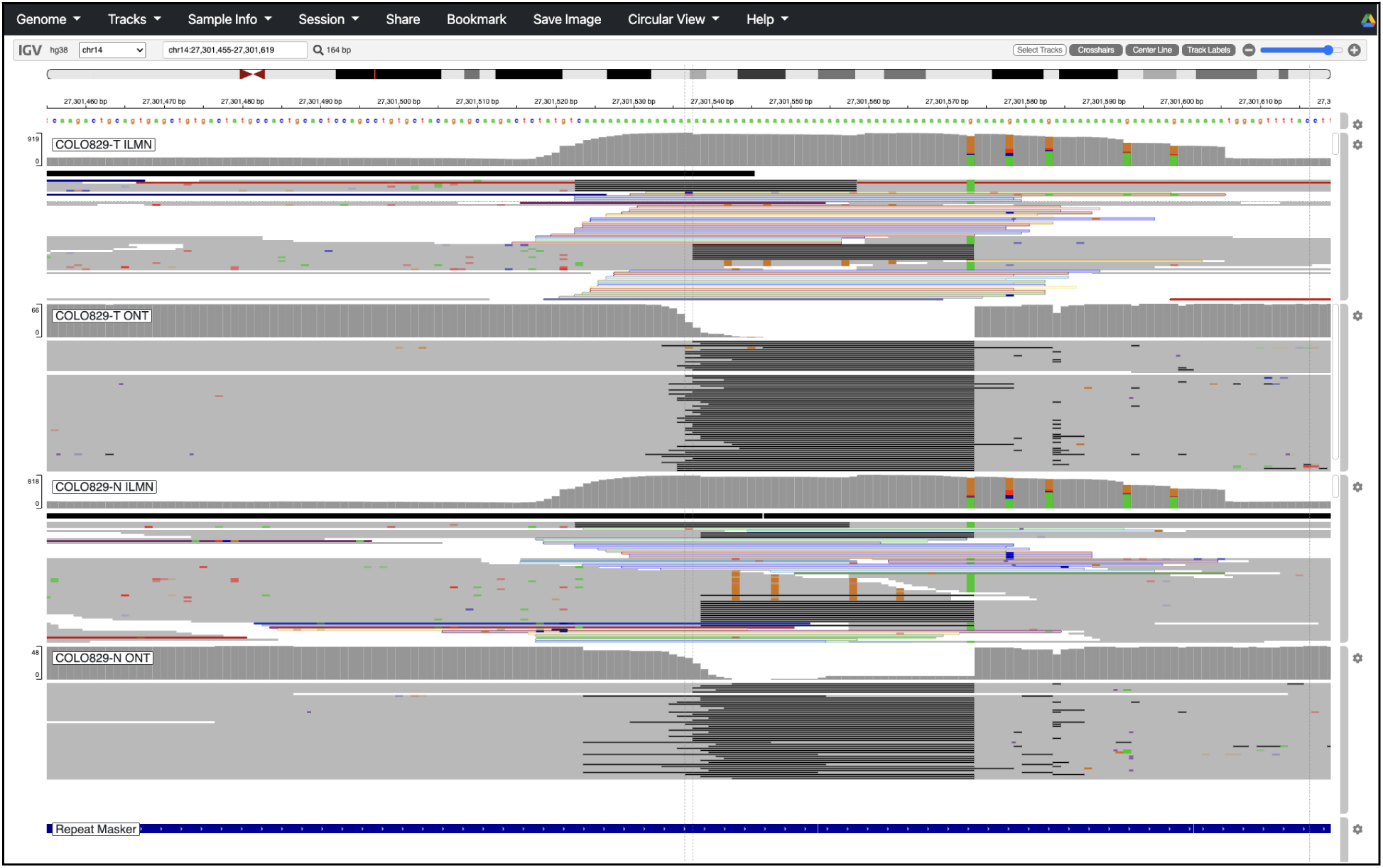
36 bp putative somatic DEL in COLO829 at chr14:27301537 dropped from the previous high confidence short read truth set. Reason – homozygous germline deletion.

**Supplementary Figure 22:**
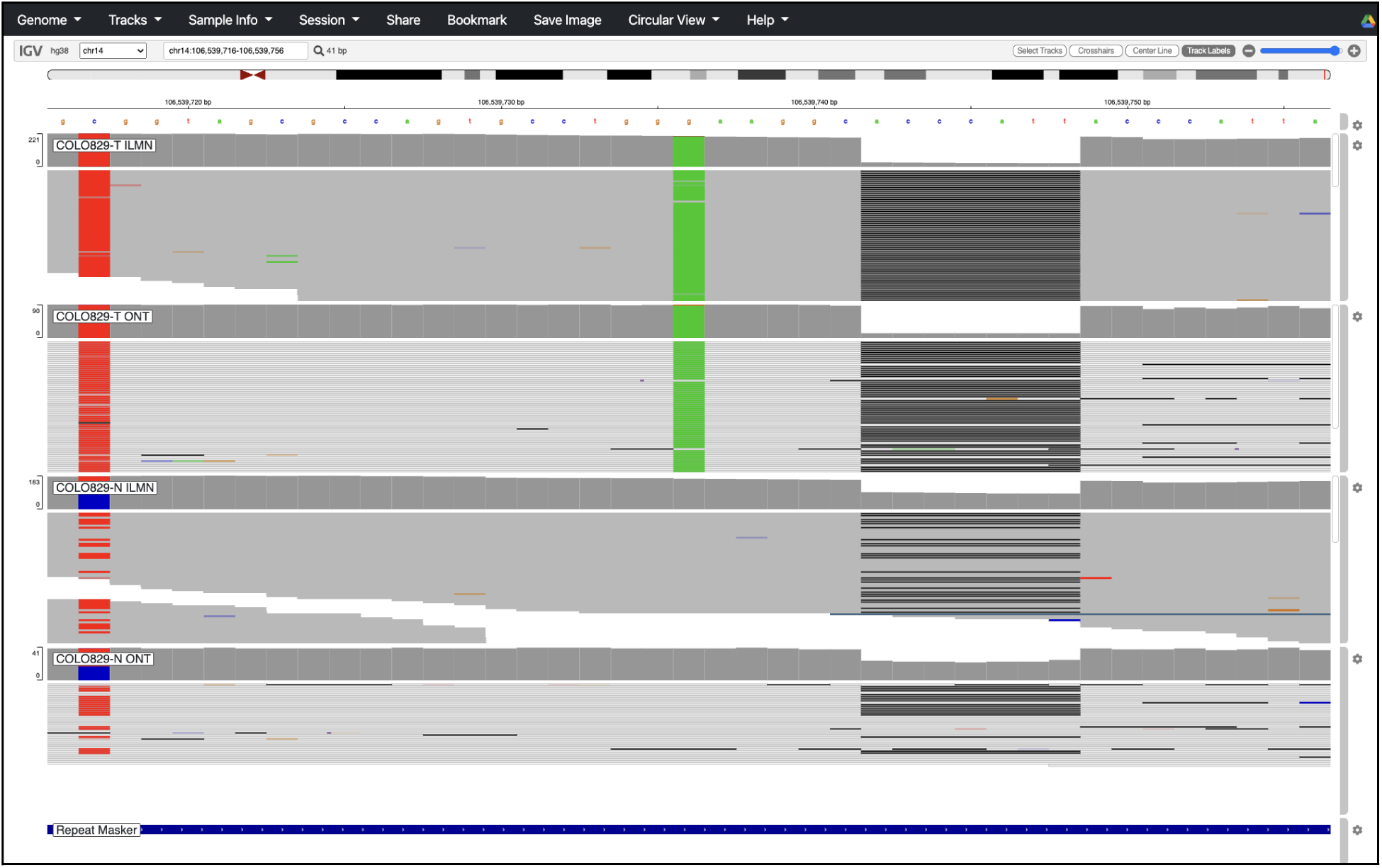
G → A putative somatic SNV in COLO829 at chr14:106539736 dropped from the previous high confidence short read truth set. Reason – only ambiguous mapping quality zero long read alignments support ALT allele.

**Supplementary Figure 23:**
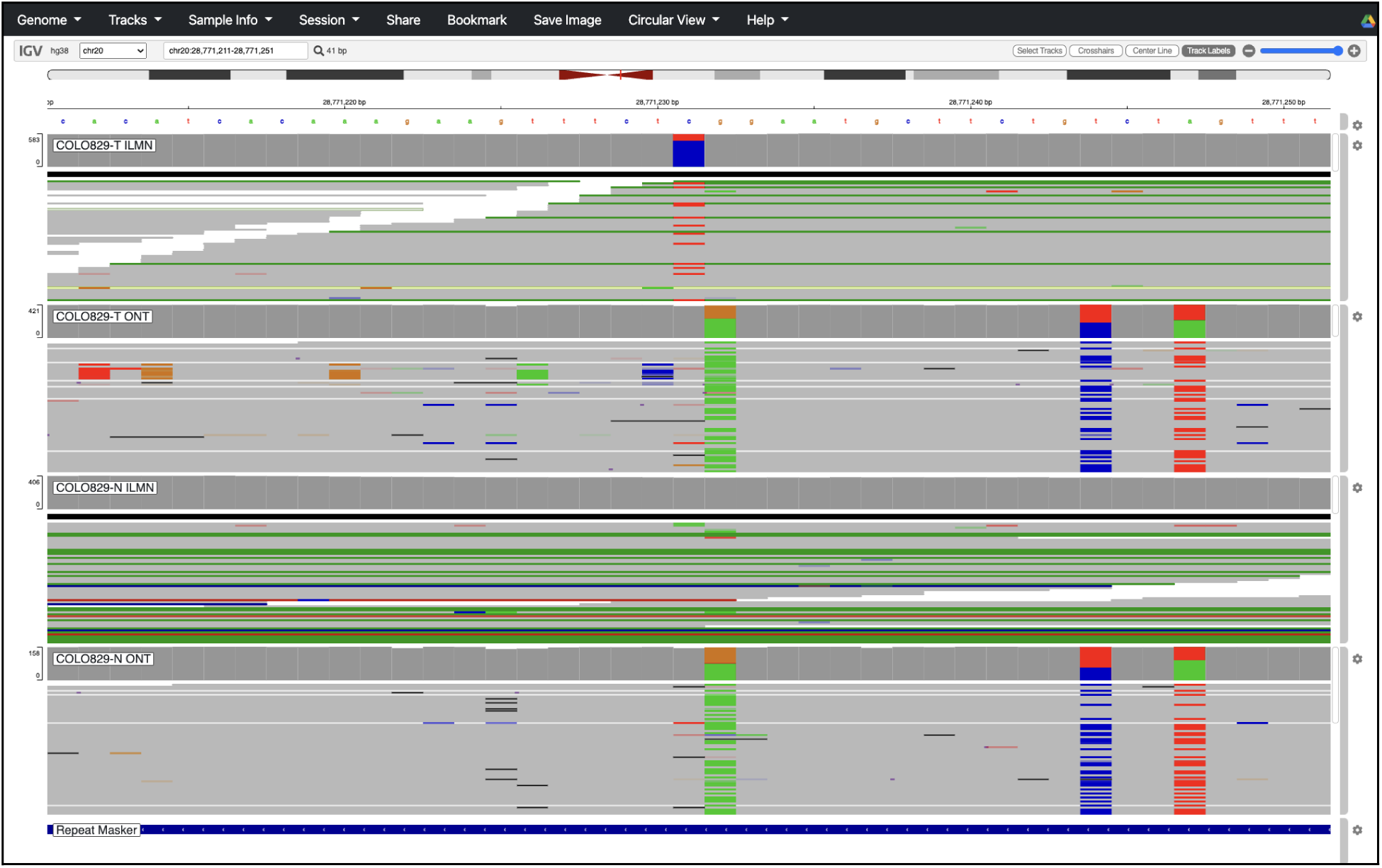
C → T putative somatic SNV in COLO829 at chr20:28771231 dropped from the previous high confidence short read truth set. Reason – ALT allele seen in normal long read alignments.

